# The evolution of recombination in self-fertilizing organisms

**DOI:** 10.1101/2022.02.24.481800

**Authors:** Roman Stetsenko, Denis Roze

## Abstract

Cytological data from flowering plants suggest that the evolution of recombination rates is affected by the mating system of organisms, as higher chiasma frequencies are often observed in self-fertilizing species compared with their outcrossing relatives. Understanding the evolutionary cause of this effect is of particular interest, as it may shed light on the selective forces favoring recombination in natural populations. While previous models showed that inbreeding may have important effects on selection for recombination, existing analytical treatments are restricted to the case of loosely linked loci and weak selfing rates, and ignore the effect of genetic interference (Hill-Robertson effect), known to be an important component of selection for recombination in randomly mating populations. In this article, we derive general expressions quantifying the stochastic and deterministic components of selection acting on a mutation affecting the genetic map length of a whole chromosome along which deleterious mutations occur, valid for arbitrary selfing rates. The results show that selfing generally increases selection for recombination caused by interference among mutations as long as selection against deleterious alleles is sufficiently weak. While interference is often the main driver of selection for recombination under tight linkage or high selfing rates, deterministic effects can play a stronger role under intermediate selfing rates and high recombination, selecting against recombination in the absence of epistasis, but favoring recombination when epistasis is negative. Individual-based simulation results indicate that our analytical model often provides accurate predictions for the strength of selection on recombination under partial selfing.

## INTRODUCTION

Genetic recombination lies at the heart of the sexual life cycle, and is often considered as one of the main evolutionary benefits of sexual reproduction (Otto, 2021). However, considerable variation for the rate and position of meiotic crossovers along chromosomes exists within and between species (Kong et al., 2010; Johnston et al., 2016; Stapley et al., 2017; Ritz et al., 2017; Brand et al., 2018; Samuk et al., 2020), showing that recombination can evolve over short timescales. Rapid changes in recombination rates have also been observed during artificial selection experiments, in response to selection on recombination itself or on other traits (reviewed in Otto and Barton, 2001). Although substantial progress has been achieved over recent years in our understanding of the molecular mechanisms governing crossover formation (Gray and Cohen, 2016; Zelkowski et al., 2019), the evolutionary forces acting on recombination in natural populations remain elusive. In most species, at least one crossover per bivalent seems required to ensure the proper segregation of chromosomes during meiosis I, while data on human trisomies suggest that homologs may fail to separate when they are entangled by too many crossovers (Koehler et al., 1996). These mechanistic constraints probably set lower and upper bounds to the genetic map length of chromosomes, but may leave some space for evolutionary change to occur. The evolution of recombination may also be affected by indirect selective forces, stemming from the effect of recombination on genetic variation. In particular, higher recombination rates may be favored when negative linkage disequilibria (LD) between selected loci exist within populations (*i.e*., when beneficial alleles at some loci tend to be associated with deleterious alleles at other loci), as recombination then increases the variance in fitness among offspring and the efficiency of natural selection (Otto and Lenormand, 2002; Agrawal, 2006). Different possible sources of negative LD have been identified, including epistatic interactions among loci (Charlesworth, 1990; Barton, 1995) and the Hill-Robertson effect, that tends to generate negative LD between selected loci from the random fluctuations of genotype frequencies occurring in finite populations (Hill and Robertson, 1966; Felsenstein, 1974; Otto and Barton, 1997; Barton and Otto, 2005; Roze and Barton, 2006). Analytical and simulation models have shown that the stochastic component of selection for recombination (due to the Hill-Robertson effect) may be stronger than deterministic components generated by epistasis even when population size is rather large, especially when linkage is tight (Otto and Barton, 2001; Keightley and Otto, 2006; Roze, 2021).

An interesting pattern observed in several genera of flowering plants is that self-fertilizing species tend to have higher chiasma frequencies than their outcrossing relatives (Roze and Lenormand, 2005; Ross-Ibarra, 2007). Detailed comparisons between the genetic maps of the selfing *Arabidopsis thaliana* and its outcrossing relative *Arabidopsis lyrata* also point to higher recombination rates in A. *thaliana* (Kuittinen et al., 2004; Hansson et al., 2006; Kawabe et al., 2006). A possible explanation for higher recombination rates in selfers could be that interhomolog polymorphism hinders recombination (Borts and Haber, 1987). However, this hypothesis does not stand up to closer scrutiny, as available data suggest that substantial levels of divergence are needed to prevent meiotic recombination (Chen and Jinks-Robertson, 1999), while moderate levels of heterozygosity may favor rather than inhibit crossovers (Benavente and Sybenga, 2004; Ziolkowski et al., 2015; Blackwell et al., 2020). Because recombination between homozygous loci has no genetic effect, selfing reduces the efficiency of recombination in breaking LD (Nordborg, 2000; Wright et al., 2008), and one may thus expect that increased rates of recombination could evolve to compensate for this effect. However, indirect selection for recombination should vanish under complete selfing (as heterozygosity should then be extremely rare), and the effect of selfing on selection for recombination may thus be non-monotonic. Furthermore, it is not immediately obvious that models for the evolution of recombination under random mating can be directly transposed to the case of partial selfers. Indeed, simulation models showed that recombination may be favored under different conditions in partially selfing than in outcrossing populations, while the exact mechanisms remained unclear (Charlesworth et al., 1977, 1979; Holsinger and Feldman, 1983).

A three-locus model on the effect of partial selfing on the evolution of recombination was analyzed by Roze and Lenormand (2005). The results showed that correlations in homozygosity across loci caused by partial selfing generate a selective force on recombination that is absent under random mating. By breaking correlations in homozygosity, recombination is favored when dominance-by-dominance epistasis is negative (meaning that double homozygotes have a lower fitness than expected based on the fitness of single homozygotes), as recombination increases the mean fitness of offspring produced by selfing. In the absence of dominance-by-dominance epistasis (or when it is positive), recombination is generally disfavored. The analysis also showed that even a small selfing rate has important consequences, the effect of breaking correlations in homozygosity quickly becoming the dominant source of indirect selection on recombination. However, the method used by Roze and Lenormand (2005) only holds when effective recombination rates are large, and thus breaks down when the selfing rate is not small, or when loci are tightly linked — yet tightly linked loci should be the ones contributing most to indirect selection for recombination, unless selfing is strong. Furthermore, this model and the previous simulation models mentioned above are deterministic, considering infinite populations. From previous results on selection for recombination in finite, randomly mating populations (Otto and Barton, 2001; Keightley and Otto, 2006; Roze, 2021), and from the fact that the effective size of highly selfing populations may be strongly reduced by interference effects among loci (Glémin and Ronfort, 2013; Roze, 2016), it seems likely that the Hill-Robertson effect should be an important component of selection for recombination in selfing organisms, but this has not been quantified.

In this article, we provide a general analysis of selection for recombination caused by interactions among deleterious alleles, in populations with arbitrary selfing rates. In a first step, we revisit Roze and Lenormand’s (2005) deterministic three locus model, and show that linkage disequilibrium is the main source of selection for recombination in the case of tightly linked loci or under strong selfing, while the effect of correlations in homozygosity stays negligible. In a second step, we explore how the stochastic component of selection for recombination (Hill-Robertson effect) is affected by the mating system, by extending Roze’s (2021) finite population model to partial selfing. Last, we extrapolate the results from the stochastic and deterministic three-locus models in order to quantify the overall strength of selection acting on a modifier allele increasing the map length of a whole chromosome, and compare the predictions obtained with results from individual-based simulations. The results confirm that the Hill-Robertson effect is often the main component of selection for recombination when the selfing rate is high, usually generating stronger benefits of recombination as selfing increases, but not always. Deterministic effects may become important under intermediate selfing rates and when the chromosomal map length is sufficiently high, and may either increase or decrease selection for recombination depending on the sign and magnitude of the different components of epistasis.

## METHODS

Our baseline model is the three-locus deterministic recombination modifier model with partial selfing considered by Roze and Lenormand (2005), that we reanalyze in order to obtain more accurate results for arbitrary selfing and recombination rates. In a second step, this model is extended to include the effect of random drift in finite populations. Finally, extrapolations are used to predict the overall strength of selection acting on a modifier affecting the genetic map length of a whole chromosome.

### The three-locus model

#### Genetic architecture

The model considers two selected loci (each with two alleles, *A, a* at the first locus and *B, b* at the second) and a recombination modifier locus (with two alleles *M* and *m*). All notations used are summarized in Table 1. Alleles *a* and b are deleterious and affect fitness by a factor 1 – *s* in homozygotes and 1 – *sh* in heterozygotes (*s* and *h* thus correspond to the selection and dominance coefficients of alleles *a* and *b*). Deleterious alleles are generated by mutation at a rate *u* per generation; back mutation is ignored. Throughout the paper, we assume that *h* is significantly greater than zero and that *u* ≪ *s*, so that the equilibrium frequency of deleterious alleles remains small. As in Roze and Lenormand (2005), epistasis between alleles *a* and *b* is decomposed into three components (see Table 2): additive-by-additive epistasis *e*_a×a_ represents the effect of the interaction between two deleterious alleles, one at each selected locus (either on the same or on different chromosomes), while additive-by-dominance epistasis *e*_a×d_ represents the effect of the interaction between three deleterious alleles, and dominance-by-dominance epistasis *e*_d×d_ the effect of the interaction between four deleterious alleles. The recombination modifier locus affects the baseline recombination rate *r_ab_* between the two selected loci, so that individuals with genotypes *MM, Mm* and *mm* at the modifier locus have recombination rates *r_ab_, r_ab_* + *δr_ab_h_m_* and *r_ab_* + *δr_ab_*, respectively, with *δr_ab_* the effect of the modifier and *h_m_* the dominance coefficient of allele *m*. Note that because recombination only affects the genotype of meiotic products when it occurs between heterozygous loci, any effect of the modifier on recombination rates between itself and the selected loci (*r_ma_, r_mb_*) will not generate any indirect selection at the modifier locus (e.g., Barton, 1995; Otto and Barton, 1997). The results given throughout the paper are valid for any ordering of the three loci along the chromosome (i.e., either *m – a – b* or *a – m – b*).

**Table 1:** Parameters and variables of the model.

|  |  |
| --- | --- |
| $\sigma$ | Selfing rate |
| $s, h, \tilde{h}$ | Selection, dominance and effective dominance coefficients of deleterious alleles |
| $e_{a \times a}, e_{a \times d}, e_{d \times d}, \tilde{e}$ | Additive-by-additive, additive-by-dominance, dominance-by-dominance epistasis and effective epistasis between deleterious alleles |
| $\beta$ | Charlesworth et al.'s (1991) synergistic epistasis coefficient |
| $\epsilon$ | Strength of selection (scaling parameter) |
| $u, U$ | Deleterious mutation rate per locus, and per chromosome |
| $r_{ij}, \tilde{r}_{ij}$ | Recombination rate and effective recombination rate between loci $i$ and $j$ |
| $\delta r_{ab}, \tilde{\delta} r_{ab}$ | Effect and effective effect of the modifier on $r_{ab}$ |
| $h_m$ | Dominance coefficient of allele $m$ at the modifier locus |
| $N, N_e$ | Census and effective population size |
| $R, R_{ES}$ | Chromosome map length and evolutionarily stable map length |
| $\delta R$ | Effect of the modifier on $R$ |
| $D_{\mathbb{U}, \mathbb{V}}$ | Genetic association between the sets $\mathbb{U}$ and $\mathbb{V}$ of loci present on the maternally and paternally inherited haplotypes of an individual |
| $\alpha_{\mathbb{U}, \mathbb{V}}$ | Effect of selection on the sets $\mathbb{U}$ and $\mathbb{V}$ of loci present on the maternally and paternally inherited haplotypes of an individual |
| $W, \bar{W}$ | Fitness of an individual and average fitness |
| $F = \sigma / (2 - \sigma)$ | Inbreeding coefficient (probability of identity by descent at one locus) |
| $\phi_{ab}$ | Joint probability of identity by descent at loci $a$ and $b$ |
| $G_{ab} = \phi_{ab} - F^2$ | Identity disequilibrium between loci $a$ and $b$ |
| $p_j, q_j$ | Frequencies of the lower- and uppercase allele at locus $j$ |
| $n$ | Mean number of deleterious alleles per chromosome |

**Table 2:** Fitness matrix in the three-locus model.

| | $AA$ | $Aa$ | $aa$ |
| --- | --- | --- | --- |
| $BB$ | 1 | $1 - hs$ | $1 - s$ |
| $Bb$ | $1 - hs$ | $(1 - hs)^2 + e_{a \times a}$ | $(1 - hs)(1 - s) + 2e_{a \times a} + e_{a \times d}$ |
| $bb$ | $1 - s$ | $(1 - hs)(1 - s) + 2e_{a \times a} + e_{a \times d}$ | $(1 - s)^2 + 4e_{a \times a} + 4e_{a \times d} + e_{d \times d}$ |

#### Genetic associations

The change in frequency at the modifier locus involves different forms of genetic associations among loci, which are defined as follows. As in Barton and Turelli (1991) and Kirkpatrick et al. (2002), we define indicator variables *X_j,ø_* and *X_ø,j_* that equal 1 if the lowercase allele at locus *j* (*m, a* or *b*) is present (or 0 if absent) on the maternally (*X_j,ø_*) or paternally (*X_ø,j_*) inherited gene of an individual. The average of these indicator variables over all individuals in the population gives the frequency of allele j on maternally and paternally inherited genes, *p_j,ø_* and *p_ø,j_*, respectively. The frequency of allele *j* in the population, *p_j_*, is thus given by (*p_j,ø_* + *p_ø,j_*)/2. Centered variables *ζ_j,ø_* and *ζ_ø,j_* are defined as:

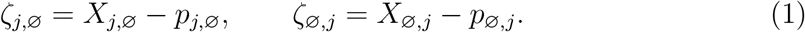

The genetic association between the sets of loci 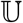 and 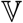 present on the maternally and paternally inherited haplotypes of the same individual is given by:

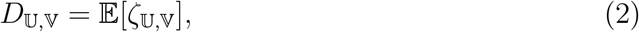

with 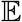 the average over all individuals in the population, and with

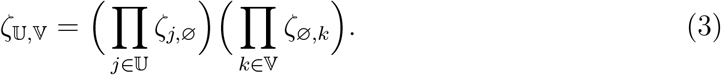

Because our model does not include and sex-of-origin effect, we always have 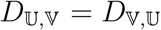. Associations between genes on the same haplotype 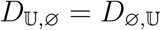 will be denoted 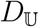 for simplicity. For example, *D_a,a_* measures the excess of homozygotes (departure from Hardy-Weinberg equilibrium) at locus *a*, while *D_ab_* is the linkage disequilibrium between alleles *a* and *b*.

#### Life cycle

Each generation starts by selection between newly formed diploid individuals, followed by meiosis and syngamy. The effect of selection on allele frequencies and genetic associations can be computed using the multilocus genetics framework of Kirkpatrick et al. (2002). For this, the fitness or an individual relative to the mean fitness of the population is written as:

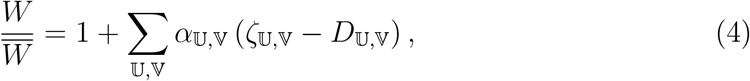

where the “selection coefficients” 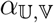 represent the effect of selection acting on the sets of loci 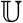 and 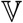 present on the maternally and paternally inherited haplotypes of an individual. Since we do not assume any sex-of-origin effect we have 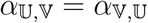, while 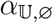 will be denoted 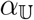 for simplicity. The combined effect of selection at loci *a* and *b* can thus be represented by 9 coefficients: *α_a_, α_b_* represent the net effect of selection against the deleterious alleles *a* and *b*, *α_a,a_, α_b,b_* the effect of dominance at the two loci, while *α_ab_, α_a,b_, α_ab,a_, α_ab,b_* and *α_ab,ab_* represent epistatic interactions, measured as deviations from additivity. Throughout the paper, we will assume that selection is weak (*s* is of order *ϵ*, where *ϵ* is a small term) while epistasis is weaker (*e*_a×a_, *e*_a×d_, *e*_d×d_ of order *ϵ*^2^). Assuming that deleterious alleles stay at low frequency, and under the fitness matrix given by Table 2, 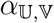 coefficients are, to leading order:

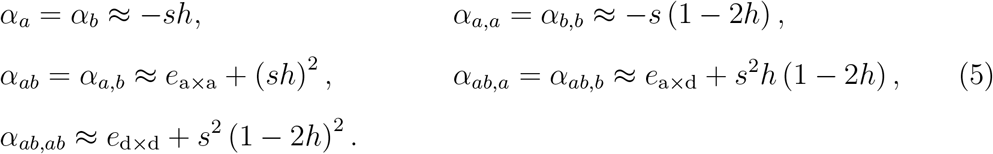

The effect of selection on genetic associations (in terms of 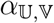 coefficients) is given by equations 9 and 15 in Kirkpatrick et al. (2002), while the change in frequency of the modifier is given by:

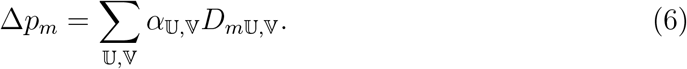

After selection, individuals produce gametes to form the zygotes of the next generation. During syngamy, a proportion *σ* of fertilizations involves two gametes produced by the same parent (selfing), while a proportion 1 – *σ* involves gametes sampled at 5 random from the whole population (outcrossing). Recombination and syngamy do not change allele frequencies, but do change genetic associations, their effect being given by equations 13 and 14 in Roze and Lenormand (2005). These equations, together with the equations describing the effect of selection on allele frequencies and genetic associations, have been implemented in a *Mathematica* notebook (available as Supplementary Material) that can be used to automatically generate recursions on these variables.

#### Approximations

Throughout the paper we assume that the effect of the recombination modifier (*δr_ab_*) is weak and compute all results to the first order in *δr_ab_*. Our three-locus diploid model can be described by 36 genotype frequencies (leading to 35 independent variables), or alternatively by 3 allele frequencies (*p_m_, p_a_, p_b_*) and 32 genetic associations. A separation of timescales argument can be used to reduce this large number of variables: in particular, when selection is weak relative to recombination, allele frequencies change slowly while genetic associations are rapidly eroded by recombination. In this case, one can show that genetic associations quickly reach a quasi-equilibrium value, which can be computed by assuming that allele frequencies remain constant (Barton and Turelli, 1991; Nagylaki, 1993; Kirkpatrick et al., 2002). In this quasi-linkage equilibrium (QLE) state, associations can be expressed in terms of allele frequencies and of the parameters of the model, and these expressions can then be plugged into equation 6 to obtain the change in frequency of the modifier. This is the approach used in Roze and Lenormand (2005) to quantify the strength of selection for recombination under partial selfing, assuming that recombination rates are sufficiently large (relative to the strength of selection) and that the selfing rate is not too large (as selfing reduces the effect of recombination). However, more accurate expressions can in principle be obtained in situations where deleterious alleles are maintained at mutation – selection balance: indeed, in this case changes in allele frequencies are only driven by the effect of the recombination modifier, and the QLE approximation thus only requires that *δr_ab_* is sufficiently small relative to effective recombination rates (Roze, 2014; Gervais and Roze, 2017; Roze, 2021). This is the approach used in the present paper to obtain approximations that remain valid under low effective recombination (i.e., when recombination rates *r_ij_* or when the outcrossing rate 1 – *σ* are of order *ϵ*).

#### The Hill-Robertson effect

An expression for the strength of selection for recombination due to the Hill-Robertson effect between two deleterious alleles in a randomly mating, diploid population was derived in Roze (2021). Generalizing this analysis to partial selfing would be extremely tedious, as a very large number of stochastic moments of genetic associations and allele frequencies would need to be computed. However, in the case of tightly linked loci (which should be the ones contributing most to selection for recombination, unless the selfing rate is very high), scaling arguments can be used to show that the effects of selection against deleterious alleles, recombination and drift under partial selfing can be predicted by replacing the dominance coefficient *h*, recombination rates *r_ij_* and the population size *N* by the effective coefficients h (1 – *F*) + *F, r_ij_* (1 – *F*) and *N/* (1 + *F*) in the expressions obtained under random mating, where *F* = *σ*/ (2 – *σ*) is the inbreeding coefficient (e.g., Nordborg, 1997; Glémin and Ronfort, 2013; Roze, 2016). We thus introduced these effective coefficients into the expression derived in Roze (2021) in order to explore how self-fertilization affects selection for recombination generated by the Hill-Robertson effect. As we will see, comparisons with multilocus simulations indicate that this approach often yields correct results.

### Multilocus extrapolation

Following Roze (2021), the three-locus analysis can be extrapolated to predict the overall strength of selection on a modifier affecting the genetic map length *R* of a whole chromosome (by an amount *δR*). For simplicity, we assume that the modifier is located at the mid-point of a linear chromosome and that the position of each crossover is sampled in a uniform distribution along the chromosome (no interference). Deleterious mutations occur at a rate *U* per haploid chromosome per generation, and we assume that all mutations have the same selection and dominance coefficients *s* and *h*, the position of each new mutation being sampled in a uniform distribution along the chromosome (infinite site model). Neglecting the effect of interactions between more than two mutations, the strength of indirect selection for recombination can be obtained by integrating the result from the three-locus analysis over the genetic map (see Supplementary Material). When the mean number of deleterious alleles per chromosome (*n*) is large and with epistasis, more accurate expressions for selection coefficients 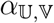 at loci *j* and *k* segregating for deleterious alleles are given by:

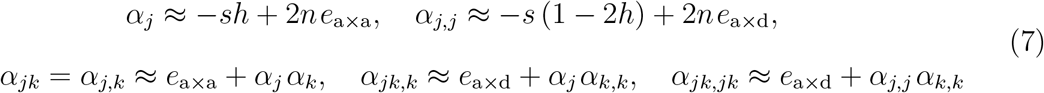

(see Supplementary Material). An approximate expression for the mean number of deleterious alleles per chromosome *n* (taking into account the effects of epistasis and selfing) can be obtained from equation 27 in Abu Awad and Roze (2020). We also computed the overall strength of selection for recombination under the diploid model of synergistic epistasis considered by Charlesworth et al. (1991), in which the fitness of individuals is given by *W* = exp [– (*αñ* + *βñ*^2^/2)], with *ñ* = *hn*_he_ + *n*_ho_ and where *n*_he_ and *n*_ho_ are the numbers of heterozygous and homozygous mutations present in the genome of the individual. As shown in Abu Awad and Roze (2020), this is equivalent to setting *e*_a×a_ = –*β h*^2^, *e*_a×d_ = –*βh* (1 – 2*h*) and *e*_d×d_ = –*β* (1 – 2*h*)^2^ in the present model.

### Individual-based simulations

Our simulation program (written in C++ and available from Zenodo) is equivalent to the program used in Roze (2021), to which partial selfing and the different forms of epistasis are added. It represents a population of *N* diploids, each carrying a pair of linear chromosomes. At each generation, the number of new deleterious mutations per chromosome is drawn from a Poisson distribution with parameter *U*, and their position along the chromosome is drawn from a uniform distribution between 0 and 1. The fitness of each individual is computed as:

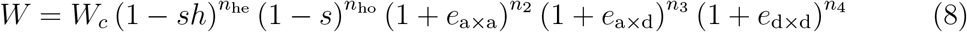

where *n*_he_ and *n*_ho_ are the number of heterozygous and homozygous deleterious mutations in the genome of the individual, and where *n*_2_, *n*_3_ and *n*_4_ are the number of interactions between 2, 3 and 4 deleterious alleles at two loci, given by:

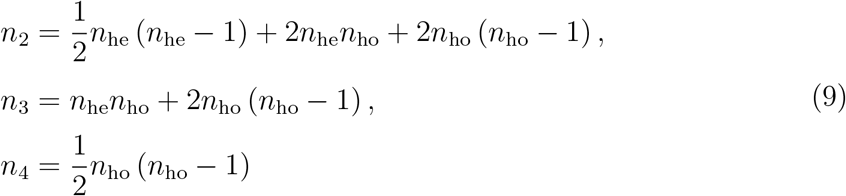

(e.g., Abu Awad and Roze, 2020). The term *W_c_* corresponds to a direct fitness effect of the chromosome map length *R*; as in Roze (2021), it is set to *W_c_* = exp (–*cR*), so that c measures a direct fitness cost per crossover. This ensures that *R* does not evolve towards very large values and allows simple comparisons with analytical predictions.

To produce individuals of the next generation, parents are sampled according to their fitness, each new individual being produced by selfing with probability *σ*. The recombination modifier locus is located at the midpoint of the chromosome, with an infinite number of possible alleles coding for different values of *R* (the map length of the individual being given by the average of its two modifier alleles). During the first 20,000 generations *R* is fixed to 1 in order to reach mutation–selection balance for deleterious alleles. Then, for an extra 5 × 10^6^ generations (increased up to 5 × 10^7^ generations for high values of *σ*), mutations are introduced at a rate 10^−4^ at the modifier locus, each mutation multiplying the value of *R* by a random number drawn from a Gaussian distribution with mean 1 and variance 0.04. To allow for large effect mutations, a proportion 0.05 of mutations have an additive effect on *R* drawn from a uniform distribution between −1 and 1 (the new value being set to zero if it is negative). The average map length 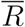, average fitness 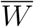, average number of deleterious mutations per chromosome *n* and number of fixed mutations are recorded every 500 generations (fixed mutations are removed from the population in order to reduce execution speed). The equilibrium value of map length is computed as the time average of 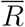, after removing the first 5 × 10^5^ generations to allow 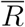 to equilibrate. In order to study the effect of variable selection coefficients of deleterious mutations, we modified the above baseline simulation program so that each new mutation is associated with a value of s drawn from a log-normal distribution (ensuring *s* > 0): the value of ln (*s*) is drawn from a Gaussian distribution with variance *sd*^2^ and mean ln 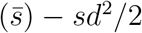 (so that the average selection coefficient 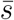 stays constant for different values of *sd*).

## RESULTS

### The deterministic three-locus model

We first reiterate the result obtained by Roze and Lenormand (2005) under high effective recombination in a more general version, and then provide a new analysis for the case of weak effective recombination. All derivations are provided in the *Mathematica* notebook available as Supplementary Material.

#### High effective recombination

As found by Roze and Lenormand (2005), the joint effects of selfing and the recombination modifier generate an association *D_mab,ab_* even in the absence of selection. This stems from the fact that recombination tends to break correlations in homozygosity between loci generated by partial selfing (see also Roze, 2009). The association *D_mab,ab_* has the sign of – *δr_ab_*, reflecting the fact that the modifier allele increasing recombination tends to be found on genetic backgrounds in which the correlation in homozygosity between loci *a* and *b* is relatively weaker. At QLE and to the first order in *δr_ab_*, it is given by:

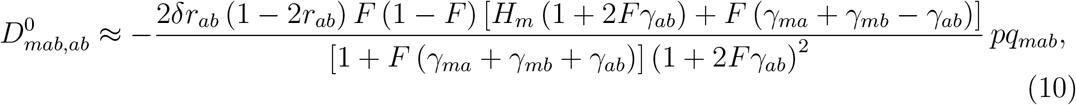

where the superscript “0” indicates that the effect of selection at loci *a* and *b* is neglected in this expression, and where *H_m_* = *h_m_* + *p_m_*(1 – 2*h_m_*), *γ_ij_* = *r_ij_*(1 – *r_ij_*) and *pq_mab_* = *p_m_q_m_p_a_q_a_p_b_q_b_* (with *q_j_* = 1 – *p_j_*). Equation 10 generalizes equation 32 in Roze and Lenormand (2005) to arbitrary *h_m_* and any ordering of the three loci along the chromosome. It shows that 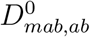 vanishes under random mating (*F* = 0) and full selfing (*F* = 1), because in these cases the mating system does not generate any correlation in homozygosity among loci.

The other associations appearing in equation 6 are generated by selection against the deleterious alleles (of order *ϵ*) and by the modifier effect. At QLE, associations *D_mi,i_, D_mij,i_* and *D_mi,ij_* (where *i, j* are either *a* or *b*) are of order *ϵ*, while associations *D_mi_, D_m,i_, D_mij_, D_m,ij_* and *D_mi,j_* are of order *ϵ*^2^ (expressions for these associations are given in Appendix A). As a result, one obtains from equations 5 and 6 that the change in frequency of the modifier is, to leading order:

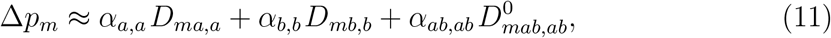

the association *D_ma,a_* being given by:

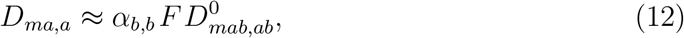

(and symmetrically for *D_mb,b_*). *D_ma,a_* is positive when *δr_αb_* > 0 and *h* < 1/2 (since *α_b,b_* and 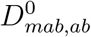 are both negative in this case), reflecting the fact that the modifier allele *m* increasing recombination tends to be found on more homozygous backgrounds at locus *a*. Indeed when *h* < 1/2, homozygous genotypes at loci a and b have, on average, a lower fitness than heterozygous genotypes, and the correlation in homozygosity generated by partial selfing increases the efficiency of selection, lowering the frequency of homozygous genotypes. As the modifier allele increasing recombination tends to break this correlation, it is associated to a relative excess in homozygosity at each selected locus. This effect disfavors recombination (since homozygotes have a lower fitness than heterozygotes on average), which is reflected by the first two terms of equation 11. Equations 11 and 12 give:

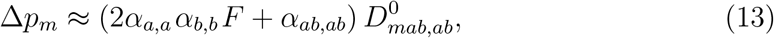

which, using equation 5, becomes:

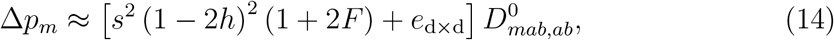

yielding Equation 36 in Roze and Lenormand (2005). Equations 10 and 14 show that increased recombination is favored when dominance-by-dominance epistasis (*e*_d×d_) is sufficiently negative. Indeed, in this case, breaking correlations in homozygosity (thus increasing the frequency of genotypes that are homozygous at one selected locus and heterozygous at the other) tends to increase the mean fitness of offspring.

#### Low effective recombination

In the previous analysis, terms in *r_mi_* (1 – *F*) appear in the denominators of the expressions for *D_mi_* and *D_m,i_* at QLE (see Appendix A), causing these expressions to diverge (i.e., tend to infinity) as effective recombination rates *r_mi_* (1 – *F*) tends to zero. Similarly, terms in *r_mab_* (1 – *F*) appear in the denominators of *D_mab_, D_m,ab_, D_ma,b_* and *D_mb,a_*, where *r_mab_* = (*r_ma_* + *r_mb_* + *r_ab_*) /2 is the probability that at least one recombination event occurs between the three loci. This indicates that these associations should play a more important role in the case of tightly linked loci. In order to explore this regime, we re-analyzed the model in the case where all recombination rates are of order *ϵ* (see Supplementary Material). This analysis shows that the effect of the linkage disequilibrium *D_ab_* between deleterious alleles (which was negligible under high effective recombination) becomes predominant when loci are tightly linked. A general expression for *D_ab_* at equilibrium in terms of 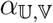 coefficients is given by:

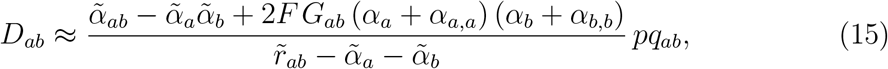

where the tilde denotes “effective coefficients”: 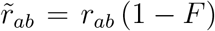, 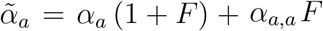 (and similarly for 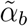), while 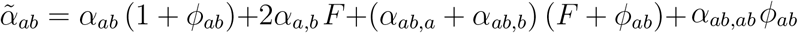, where *ϕ_ab_* is the probability of joint identity-by-descent at the two loci (which is approximately *F* when *r_ab_* is small, see Appendix A). Finally, *G_ab_* in equation 15 refers to the identity disequilibrium between the two loci, defined as *G_ab_* = *ϕ_ab_* – *F*^2^ (Weir and Cockerham, 1973), and thus approximately equal to *F* (1 – *F*) under tight linkage. Equation 15 simplifies to (*α_ab_* – *α_a_α_b_*) *pq_ab_*/ (*r_ab_* – *α_a_* – *α_b_*) in the absence of selfing; this is equivalent to the result obtained by Barton (1995) under strong recombination (equation 9b in Barton, 1995), except that *α_a_* and *α_b_* now appear in the denominator, due to our assumption that *r_ab_* is small (of order *ϵ*). With selfing, two important differences appear: (*i*): the recombination rate and selection coefficients are replaced by effective coefficients 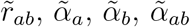 (since increased homozygosity affects both the effects of recombination and of selection, as explained in Appendix B), and (*ii*): an extra term, involving the identity disequilibrium *G_ab_*, appears in the numerator and tends to generate positive linkage disequilibrium between deleterious alleles.

Using the expressions for 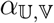 coefficients given by equation 5, equation 15 simplifies to:

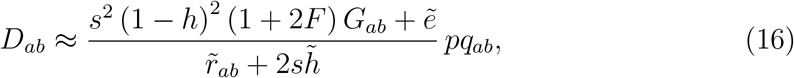

with 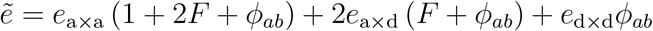 and 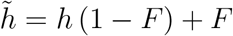. The fact that partial selfing generates positive linkage disequilibrium between deleterious alleles in the absence of epistasis has been noticed in previous analytical and simulation studies (Roze and Lenormand, 2005; Kamran-Disfani and Agrawal, 2014), and may be understood as follows. Lineages with different histories of inbreeding coexist in a partially selfing population: lineages that have been inbred for many generations tend to be very homozygous (at all loci), while lineages that have been inbred for fewer generations tend to be less homozygous. This is the basis of correlations in homozygosity among loci, represented by the identity disequilibrium *G_ab_*. Because homozygosity increases the efficiency of selection, the frequency of deleterious alleles tends to be lower within lineages that have been inbred for longer (purging), and higher in less inbred lineages, resulting in positive linkage disequilibrium between deleterious alleles. As shown by equation 16, *D_ab_* may become negative when epistasis is negative and sufficiently strong, the relative importance of *e*_a×d_ and *e*_d×d_ increasing as the selfing rate increases.

The fact that the allele coding for higher recombination tends to erode *D_ab_* more rapidly in turn generates genetic associations between the modifier and the selected loci. In particular, we have:

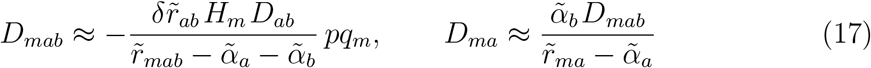

with 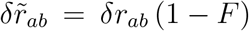, 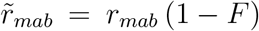, and again 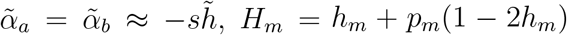 (*D_mb_* is given by a symmetric expression). These equations take the same form as under random mating (e.g., equations 9c and 11 in Barton, 1995) and are interpreted in the same way: when *D_ab_* > 0, *AB* and *ab* haplotypes are in excess in the population, and the allele increasing recombination (allele *m* if *δr_ab_* > 0) tends to reduce this excess, and thus becomes more associated with *Ab, aB* haplotypes (which is reflected by a negative value of *D_mab_*). By contrast, when *D_ab_* < 0, the allele increasing recombination becomes more associated with *ab, AB* haplotypes (which is reflected by a positive value of *D_mab_*). When *D_mab_* > 0, selection is more efficient in the *m* background (because the frequency of extreme haplotypes *AB* and *ab* is higher in this background), and *m* thus becomes better purged from deleterious alleles (which is reflected by negative values of *D_ma_, D_mb_*). When *D_mab_* < 0, selection is less efficient in the *m* background, leading to positive values of *D_ma_, D_mb_* (deleterious alleles are more frequent in the *m* background).

Under weak recombination, separation of timescale arguments can be used to express the other associations that appear in equation 6 in terms of *D_mab_, D_ma_* and *D_mb_* (e.g., Roze, 2016); one obtains:

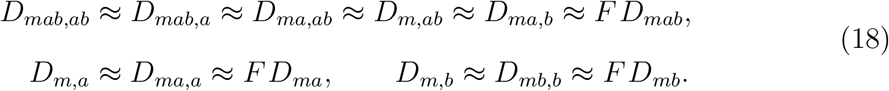

Plugging these into equation 6 yields:

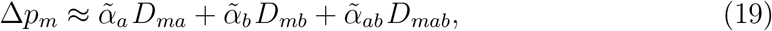

which again takes the same form as under random mating (equation 8 in Barton, 1995).

From equations 17 and 19, one obtains:

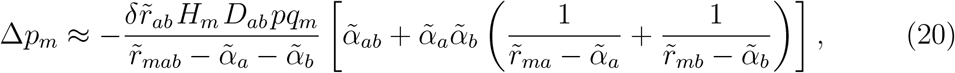

equivalent to equation 12 in Barton (1995). Given that the equilibrium frequency of deleterious alleles is approximately 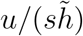, thus of order *u/ϵ*, equations 16 and 20 show that under weak effective recombination (i.e., when 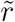 coefficients are of order *ϵ*), selection for recombination generated by the linkage disequilibrium *D_ab_* is of order *δr_ab_u*^2^/*ϵ*, thus stronger than selection for recombination generated by the term in 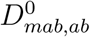 seen in the previous subsection (of order *δr_ab_u*^2^, see equation 14). When 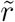 coefficients are not small (of order 1), however, indirect selection generated by *D_ab_* becomes of order *δr_ab_u*^2^*ϵ*^2^ (and thus negligible). A general expression for the change in frequency of the modifier, valid under both weak and strong effective recombination, can thus be obtained by summing equations 14 and 20. We noticed that in some cases, more accurate expressions can be obtained by taking into account indirect effects of 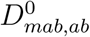 on other genetic associations between the modifier and selected loci (these terms should be negligible under both weak and strong effective recombination, but improve the precision of our approximations for intermediate values of recombination rates); these expressions are given in Appendix C. Two further improvements to our approximations are done in Appendix C, leading to more accurate expressions when epistasis is of the same order of magnitude as the strength of selection (*e*_a×a_, *e*_a×d_, *e*_d×d_ of order *ϵ*) and when the selfing rate is high (see Supplementary Material for derivations).

Figures 1 and S1 show that our approximations provide correct predictions for the change in frequency at the modifier locus, for all values of the selfing rate between 0 and 1. The dots in these figures correspond to the results of deterministic simulations, obtained by iterating exact recurrence equations for the 36 genotype frequencies: allele *M* is fixed during the first 3000 generations to reach mutation–selection balance at the selected loci, then allele *m* is introduced in frequency 0.01 and the population is let to evolve for an extra 1000 generations, Δ*p_m_*/*pq_mab_* being averaged over the last 500 generations (see Supplementary Material). As can be seen on Figure 1, indirect selection is mostly driven by *D_ab_* when recombination rates are small (left figures) or when the selfing rate approaches 1. In the absence of epistasis, the identity disequilibrium generates positive *D_ab_* which disfavors recombination (as breaking positive *D_ab_* reduces the variance in fitness, see Figures 1A, 1B). Negative epistasis generates negative *D_ab_*, which favors increased recombination when effective recombination rates are sufficiently small (equation 20, Figures 1C-F). The relative effect of the term in 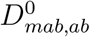 becomes more important in the case of loosely linked loci and for intermediate selfing rates, either when epistasis is absent (in which case it disfavors recombination, see equation 14, Figure 1B), or when the dominance-by-dominance component of epistasis is important (in which case it favors increased recombination when *e*_d×d_ < 0, Figure 1F). Finally, selection on recombination vanishes under complete selfing (*σ* = 1), as recombination is ineffective in that case.

**Figure 1.**
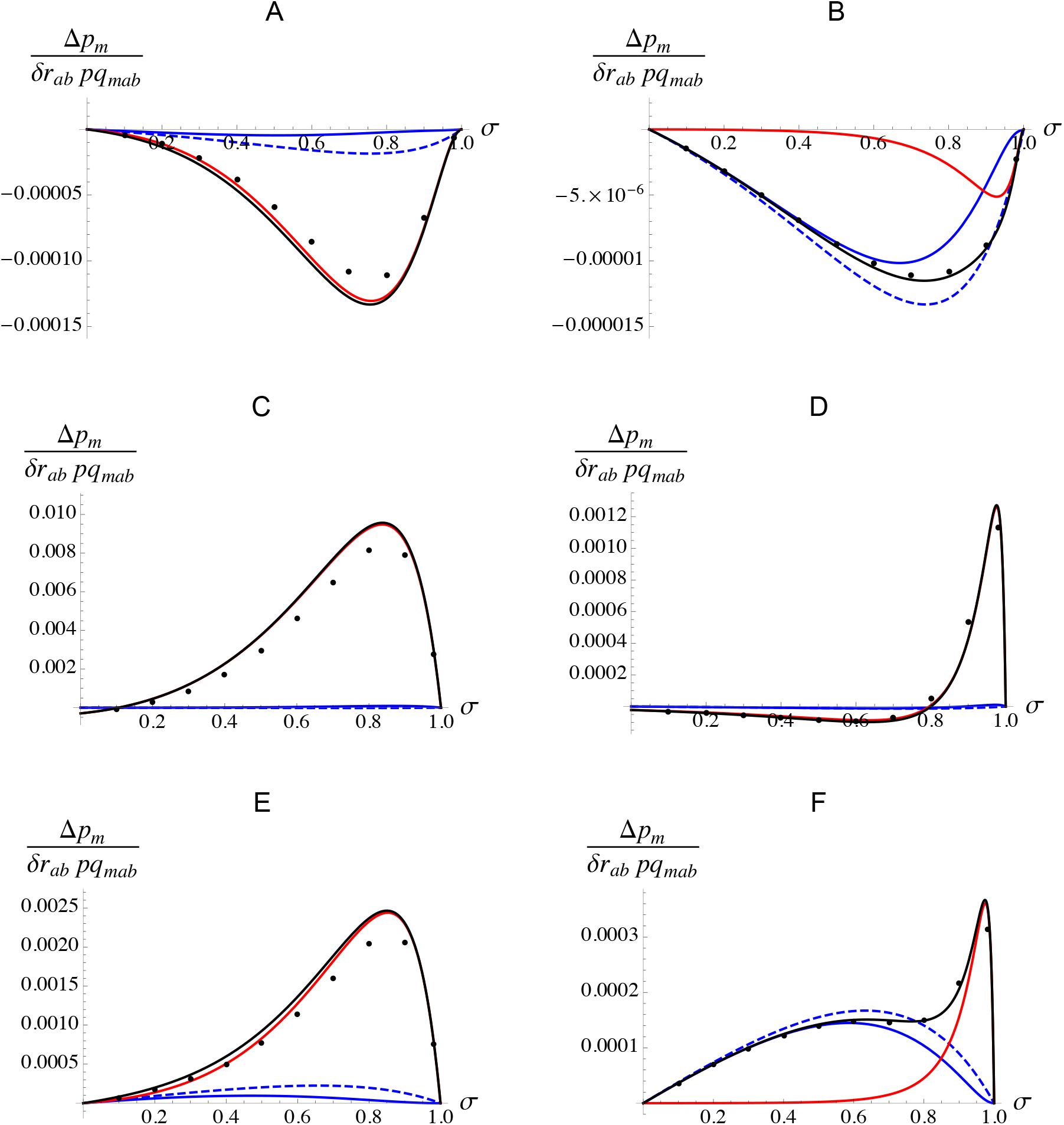
Effect of the selfing rate *σ* on the change in frequency of allele *m* in the deterministic three-locus model (scaled by *δr_ab_ pq_mab_*) with no epistasis (A and B), negative additive-by-additive epistasis (*e*_a×a_ = –0.001; C and D) and negative dominance-by-dominance epistasis (*e*_d×d_ = –0.001; E and F), and for different recombination rates (A, C, E: *r_ma_* = *r_ab_* = 0.01; B, D, F: *r_ma_* = *r_ab_* = 0.1). Dots correspond to deterministic simulation results (see text for details) and black curves to the result obtained from equations 6 and C1 – C11, which is the sum of a term generated by *D_ab_* – *D_ab_* (red curves) and a term generated by 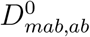 (blue curves); dashed blue curves correspond to the strong recombination approximation (Equation 11). Note that the red and black curves are nearly undistinguishable in C, D. The effect of additive-by-dominance epistasis (*e*_a×d_) is similar to the effect of additive-by-additive epistasis (*e*_a×a_) for these parameter values, and is shown in Figure S1. Loci are in the order *m* – *a* – *b*, parameters values are: *s* = 0.01, *h* = 0.2, *h_m_* = 0.5, while *δr_ab_* = 0.01 and *u* = 10^−5^ in the simulations.

### The Hill-Robertson effect

An expression for the strength of selection for recombination caused by the Hill-Robertson effect between two deleterious alleles in a diploid, randomly mating population was derived by Roze (2021). This expression consists in a sum of terms corresponding to different mechanisms generating selection for recombination, that all take the form 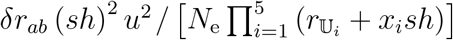 where *N*_e_ is the effective population size, 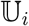 is either *mab, ma, mb* or *ab*, and *x_i_* equals 1, 2, 3 or 4. In the case of loosely linked loci 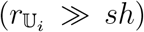, the terms in *sh* in the denominator may be neglected and the strength of selection for recombination increases as *sh* increases — being roughly proportional to (*sh*)^2^. In the case of tightly linked loci 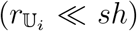, however, recombination rates 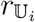 may be neglected and selection for recombination now decreases as sh increases — being roughly proportional to 1/ (*sh*)^3^.

In order to extend these results to partially selfing populations, we replaced *δr_ab_*, 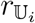 and *h* by the effective parameters 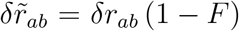, 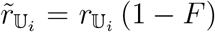 and 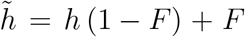 in the expression derived in Roze (2021). The effect of increasing selfing (thus increasing *F*) on the strength of selection for recombination due to the Hill-Robertson effect can be understood using the same reasoning as above. When effective recombination rates are large relative to the strength of selection against deleterious alleles (selfing rate not too large, 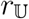 not too small so that 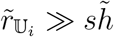), selection for recombination becomes approximately proportional to 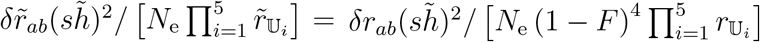, which increases as *F* increases (mostly due to the decreased effective recombination rates 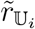 at the denominator, but also to the increase in 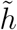). When effective recombination rates are small relative to the strength of selection (high selfing and/or tightly linked loci, so that 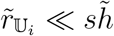), selection for recombination is approximately proportional to 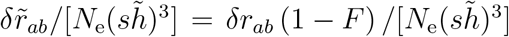, which decreases as *F* increases (mostly due to the decreased effect of the modifier 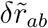, but also to the increase in 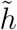). One thus predicts that increasing selfing should generally have a non-monotonic effect on selection for recombination due to the Hill-Robertson effect (selection for recombination first increasing, and then decreasing towards zero as *F* increases from 0 to 1), the position of the maximum depending on *sh* and on the values of recombination rates (however, selfing may always decrease selection for recombination when *sh* is sufficiently strong and/or linkage sufficiently tight). These verbal predictions are illustrated by Figure 2. Note that selfing also has the additional effect of reducing *N*_e_ by a factor 1/ (1 + *F*) (Pollak, 1987; Nordborg, 2000), thus increasing the strength of the Hill-Robertson effect, but this effect stays minor relative to the effect of selfing on effective recombination rates. Selfing may cause stronger reductions in *N*_e_ when deleterious alleles are segregating at many linked loci, however, through background selection effects (Glémin and Ronfort, 2013; Roze, 2016).

**Figure 2.**
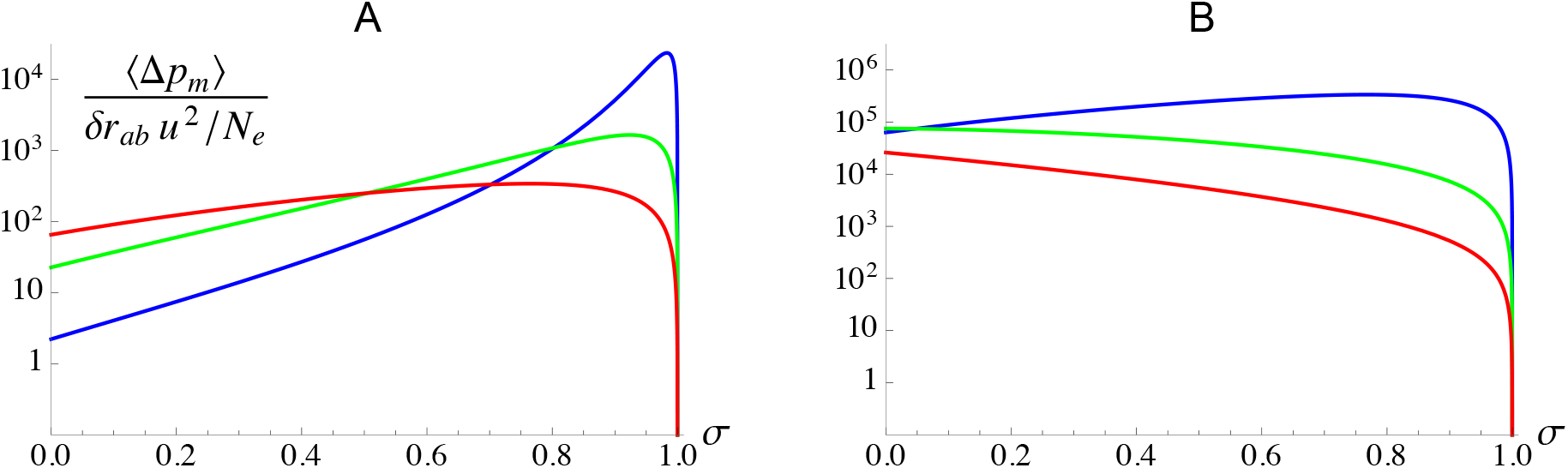
Effect of the selfing rate σ on the expected change in frequency of allele *m* (scaled by *δr _ab_ u*^2^/*N*_e_ and on a log scale) generated by the Hill-Robertson effect between alleles *a* and *b*, for different values of the strength of selection against deleterious alleles (blue: *s* = 0.005; green: *s* = 0.02; red: *s* = 0.05) and recombination rates (A: *r_ma_* = *r_ab_* = 0.05; B: *r_ma_* = *r_ab_* = 0.005). The value of *σ* maximizing selection for recombination decreases as *s* increases and as recombination decreases. When selection is sufficiently strong and linkage sufficiently tight, selection for recombination decreases monotonously as *σ* increases (as can in seen in B for *s* = 0.02, *s* = 0.05). Loci are in the order *m* – *a* – *b*, the dominance coefficient of deleterious alleles is set to *h* = 0.2.

These results also provide us with some understanding of the relative importance of stochastic and deterministic sources of selection for recombination: when effective recombination rates are large, indirect selection due to the Hill-Robertson effect is of order *δr_ab_u*^2^*ϵ*^2^/*N*_e_, and should thus be negligible relative to the deterministic component (of order *δr_ab_u*^2^). When effective recombination rates are small (of order *ϵ*), however, selection due to the Hill-Robertson effect is now of order *δr_ab_u*^2^/(*N*_e_*ϵ*^3^), and may thus become of the same order of magnitude or stronger than deterministic terms (which are then of order *δr_ab_u*^2^/*ϵ*, as shown in the previous subsection).

### Multilocus extrapolation

The three-locus analysis can be extended to compute the overall strength of indirect selection acting on a modifier affecting the genetic map length of a whole chromosome. Neglecting the effect of interactions between deleterious alleles at more than two loci, this can be done by integrating the result from the three-locus model over all possible positions of alleles *a* and *b*. As in Roze (2021), we consider a linear chromosome with map length R (in Morgans), along which deleterious mutations occur at a rate *U* per chromosome per generation. The modifier is located at the mid-point of the chromosome, allele m increasing map length by an amount *δR*/2 when heterozygous and *δR* when homozygous (see Methods). The strength of selection for allele *m*, defined as 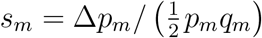, can be decomposed into three terms:

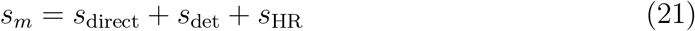

where *s*_direct_ corresponds to the effect of direct selective pressures acting on *R* (due to any direct effect of *R* on the fitness of individuals), *s*_det_ to indirect selection caused by deterministic interactions between deleterious alleles, and *s*_HR_ to indirect selection caused by the Hill-Robertson effect. Assuming a direct fitness cost of crossovers so that fitness decreases as *e^−cR^* as *R* increases, the direct selection term is given by *s*_direct_ = –*cδR* (1 + *F*) to the first order in *δR* (e.g., Gervais and Roze, 2017). From the analysis above, the term *s*_det_ can be further decomposed into 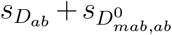, where *s_D_ab__* corresponds to the overall effect of deterministically generated linkage disequilibria (*D_ab_*) between deleterious alleles (either by epistasis or by identity disequilibria), given by equations 15 and 20, while 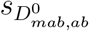 corresponds to the overall effect of associations of the form *D_mab,ab_* generated by correlations in homozygosity and by the modifier effect, given by equations 10 and 13. Because *s_D_ab__* should be mostly driven by tightly linked loci, recombination rates are approximated by genetic distances in equations 15 and 20 (before integrating over the genetic map), while *δr_ab_* is approximated by *δRr_ab_*/*R* (Roze, 2021). By contrast, loosely linked loci can make a stronger contribution to 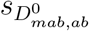, and the recombination rate between loci *i* and *j* in equation 10 is thus expressed in terms of the genetic distance *d_ij_* between these loci using Haldane’s mapping function *r_ij_* = [1 – > *e*^−2*d_ij_*^]/2 (Haldane, 1919), while *δr_ab_* is approximated by *δRe*^−2*d_ab_*^ *d_ab_/R* (see Supplementary Material). Note that when *U* is not small (so that the mean number of deleterious alleles per chromosome *n* may be large), and with epistasis, the expressions for 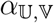 coefficients given by equation 7 must be used instead of equation 5.

Concerning the integration of the term generated by the Hill-Robertson effect, Roze (2021) showed that using scaled recombination rates 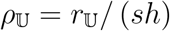 eliminates *sh* from the integrand, *sh* only appearing in the integration limits, given by *R*/ (2*sh*). The same method can be used when 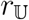, *δr_ab_* and *h* are changed to effective parameters in order to incorporate the effect of selfing. In particular, defining scaled recombination rates 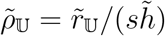, one obtains that the overall strength of indirect selection generated by the Hill-Robertson effect is given by the same expression as under random mating (equation 3 in Roze, 2021), except that the integration limits become 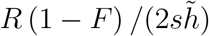 and that the whole expression is multiplied by a factor 1/ (1 – *F*)^2^. When 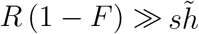 (which implies that the selfing rate is not too large and *s* is sufficiently small), the integral can be approximated by the same integral taken between 0 and infinity, which is ≈ 1.8 (Roze, 2021), yielding:

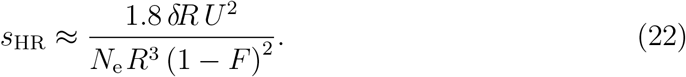

Equation 22 is equivalent to equation 1 in Roze (2021), replacing *R* by 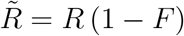 and *δR* by 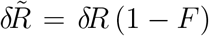. Note that this approximation is not valid when *F* approaches 1, however, as 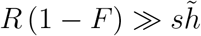 cannot hold in this case. Finally, the effective population size *N*_e_ may be significantly lowered by background selection effects when *U* is not small. Assuming that the reduction in *N*_e_ is mostly due to tightly linked loci, one obtains that 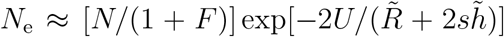 (e.g., Hudson and Kaplan, 1995; Roze, 2016), where *N* is the census population size.

The evolutionarily stable map length (*R*_ES_) corresponds to the value of *R* for which direct and indirect selection balance each other, that is, *s_m_* = 0. Figure 3 shows *R*_ES_ as a function of the selfing rate *σ*, for different values of the direct cost of recombination c. The curves correspond to the analytical predictions, obtained using *Mathematica* by numerically integrating the results of the three-locus model over the genetic map to obtain *s*_det_ and *s*_HR_ for a range of values of R, and finding the value of *R* for which *s_m_* = 0 by interpolation (see Supplementary Material). Note that the more precise approximations given in Appendix C have been used to compute *s*_det_, but using equations 10, 13, 15 and 20 often yields similar results. Figure 3 shows that the ES map length generally increases as the selfing rate increases, due to stronger indirect selection caused by the Hill-Robertson effect. While indirect selection vanishes under full selfing (*σ* = 1), the analytical model predicts that the maximum map length is reached for values of *σ* very close to 1, in particular when *c* = 10^−3^ and *c* = 10^−4^. In the absence of epistasis, the deterministic component of indirect selection selects against recombination (*s*_det_ < 0). Figure 3 shows that this deterministic component stays negligible relative to the Hill-Robertson effect for parameter values leading to low *R*_ES_ (*c* = 0.01), while its relative effect becomes more important for parameter values leading to higher *R*_ES_ (*c* = 10^−4^). This agrees with the prediction that the Hill-Robertson effect becomes stronger than deterministic effects when effective recombination rates are sufficiently small. In the absence of epistasis, we generally found that *s*_det_ is mainly driven by *s_D_ab__*, the effect of 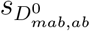 staying negligible (not shown). Figure 3 also shows that extrapolations from our three-locus model often provide accurate predictions of the evolutionarily stable map length observed in the simulations, discrepancies appearing for high values of the selfing rate. These discrepancies may be due to the fact that using effective recombination coefficients to transpose the result obtained under random mating to the case of a partially selfing population (as we did to compute *s*_HR_) should strictly only hold when loci are sufficiently tightly linked (e.g., Padhukasahasram et al., 2008; Roze, 2016), while loosely linked loci may significantly contribute to selection for recombination when the selfing rate is high.

**Figure 3.**
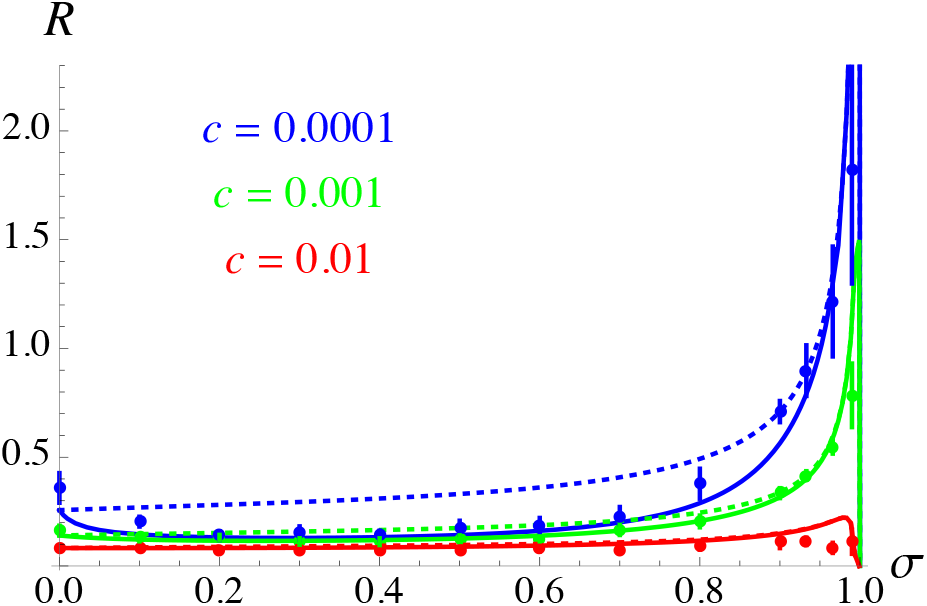
Effect of the selfing rate *σ* on the evolutionarily stable map length *R*_ES_, for different values of the cost of recombination c. Solid curves correspond to the analytical predictions obtained by solving *s*_direct_ + *s*_det_ + *s*_HR_ = 0 for *R*, where *s*_det_ and *s*_HR_ are obtained by integrating the expressions for the strength of indirect selection generated by deterministic effects (for *s*_det_) and by the Hill-Robertson effect (for *s*_HR_) over the genetic map (see Supplementary Material). Dotted curves correspond to the predictions obtained when ignoring indirect selection caused by deterministic effects (i.e., solving *s*_direct_ + *s*_HR_ = 0 for *R*). Dots correspond to individual-based simulation results; in this and the following figures, error bars are obtained by dividing the simulation output (after removing the first 5 × 10^5^ generations) into 10 batches and calculating the variance of the average map length per batch, error bars measuring ±1.96 SE. Parameter values are *N* = 20,000, *U* = 0.1, *s* = 0.02, *h* = 0.2, *e*_a×a_ = *e*_a×d_ = *e*_d×d_ = 0.

Figure 4 shows the effects of the deleterious mutation rate *U*, population size *N*, selection and dominance coefficients of deleterious alleles (*s, h*) on the evolutionarily stable map length (Figure S2 shows the same results with *R* on a log scale). As predicted, increasing U and/or decreasing *N* leads to stronger effects of Hill-Robertson interference between deleterious alleles, favoring higher values of *R*. As can be seen on Figure 4A, our analytical approximations overestimate the ES map length when *U* is high: this is probably due to the effect of higher-order associations (involving more than two selected loci) that are not taken into account in the analysis. Furthermore, in some simulations with *U* = 0.5, *R* fell down to a very small value at some point during the simulation, which led to a quick accumulation of heterozygous mutations, the only surviving individuals being heterozygous for haplotypes carrying different deleterious alleles in repulsion (pseudo-overdominance, e.g., Charlesworth and Charlesworth, 1997; Pálsson and Pamilo, 1999; Waller, 2021). This occurred for 0.5 ≤ *σ* ≤ 0.7, in which case the simulation had to be stopped manually, the number of segregating mutations quickly becoming very large. Figure 4C shows that selection for recombination is stronger when deleterious alleles are more weakly selected, as already found by Roze (2021); furthermore, above a given value of s, the equilibrium map length decreases as the selfing rate increases (which can be understood from the reasoning given in the previous subsection and Figure 2). By contrast, the dominance coefficient *h* of deleterious alleles has only little effect on the ES map length (Figure 4D). Figures S3 and S4 show that the results are not significantly affected by introducing variability in the selection coefficients of deleterious alleles into the simulation program, nor by limiting to 100 or 1000 the number of loci at which deleterious mutations may occur. Figure S5 shows that the strength of selection for recombination due to the Hill-Robertson effect is accurately predicted by equation 22 when selection against deleterious alleles is sufficiently weak, as long as the selfing rate is not too high.

**Figure 4.**
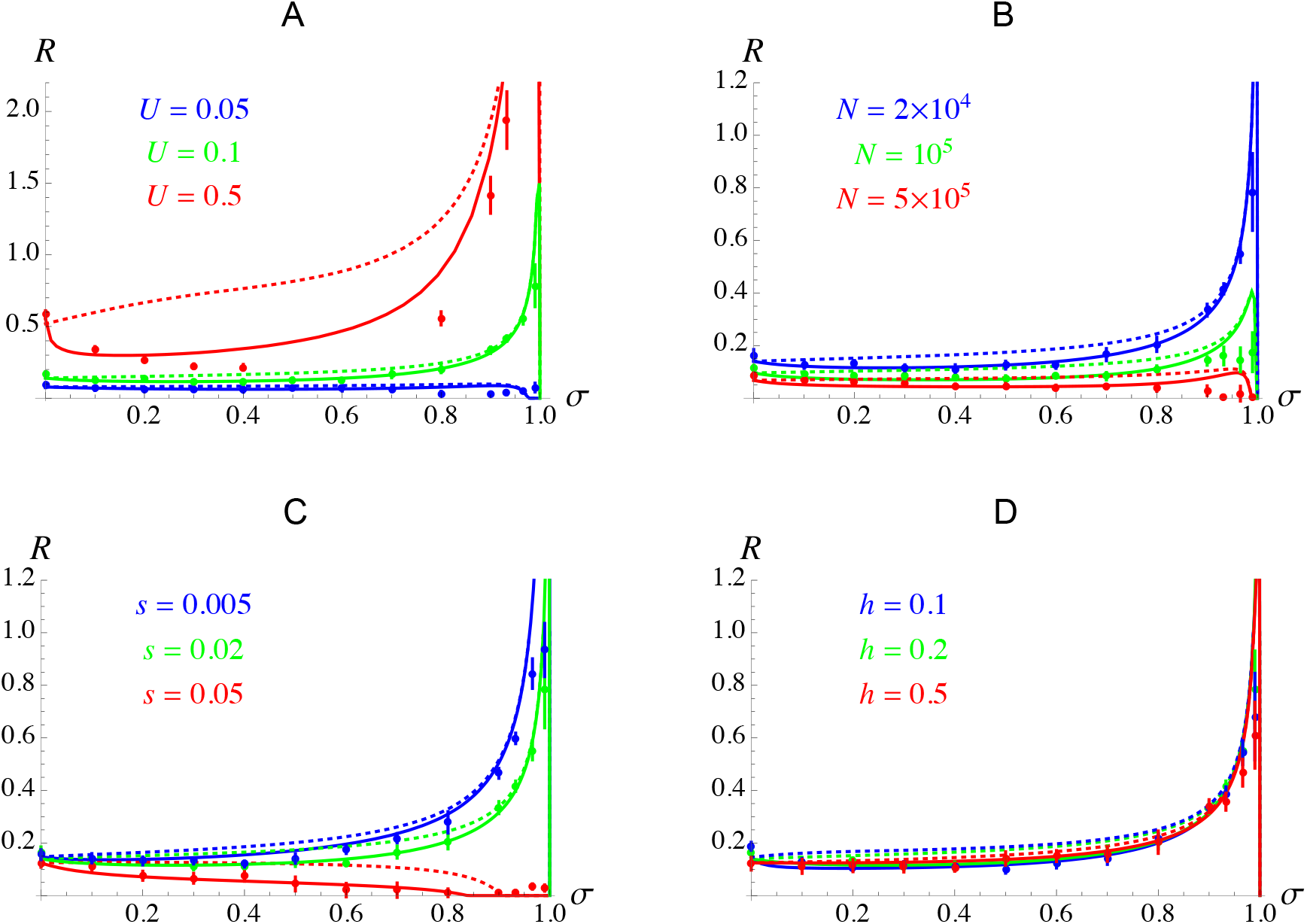
Effect of the selfing rate σ on the evolutionarily stable map length *R*_ES_, for different values of the mutation rate *U*, population size *N*, strength of selection and dominance coefficient of deleterious alleles (*s, h*). Dots correspond to simulation results, curves have the same meaning as in Figure 3, and default parameter values are as in Figure 3 with *c* = 0.001 (no epistasis).

Figure 5 shows the effect of each form of diploid epistasis (*e*_a×a_, *e*_a×d_, *e*_d×d_) and of Charlesworth et al.’s (1991) synergistic epistasis coefficient *β* (that combines the three forms of epistasis, see Methods) on the ES map length. Note that only negative values of epistasis were considered, as combinations of deleterious mutations quickly become beneficial when epistasis is positive and *U* is not very small. Negative epistasis tends to lower the strength of the Hill-Robertson effect (dotted curves on Figure 5), by increasing the effective strength of selection against deleterious alleles. Despite that, the deterministic effects generated by negative epistasis increase the overall strength of selection for recombination. Figure S6 shows that this increase is driven by the term in *D_ab_* in the case of negative additive-by-additive (*e*_a×a_) and additive-by-dominance (*e*_a×d_) epistasis, the term in 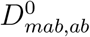 being negligible when *e*_a×a_ < 0, while it disfavors recombination when *e*_a×d_ < 0 (one can show that this last effect is generated by the first term of equation 13, since effective dominance coefficients *a_j,j_* are increased by *e*_a×d_, as shown by equation 7). In the case of dominance-by-dominance epistasis (*e*_d×d_), the important increase in recombination observed for moderate values of the selfing rate is generated by the term in 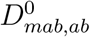, and thus corresponds to the benefit of recombination previously described by Roze and Lenormand (2005). One can note that the model overestimates the strength of selection for recombination in this case. This is possibly due to the fact that, while the effect of *e*_d×d_ on effective dominance coefficients *j* is negligible as long as epistasis is sufficiently weak, one can show that *a_j,j_* increases with *e*_d×d_ when dominance-by-dominance epistasis becomes the main source of selection against deleterious alleles (see Appendix D in Roze, 2009), reducing selection for recombination through the first term of equation 13.

**Figure 5.**
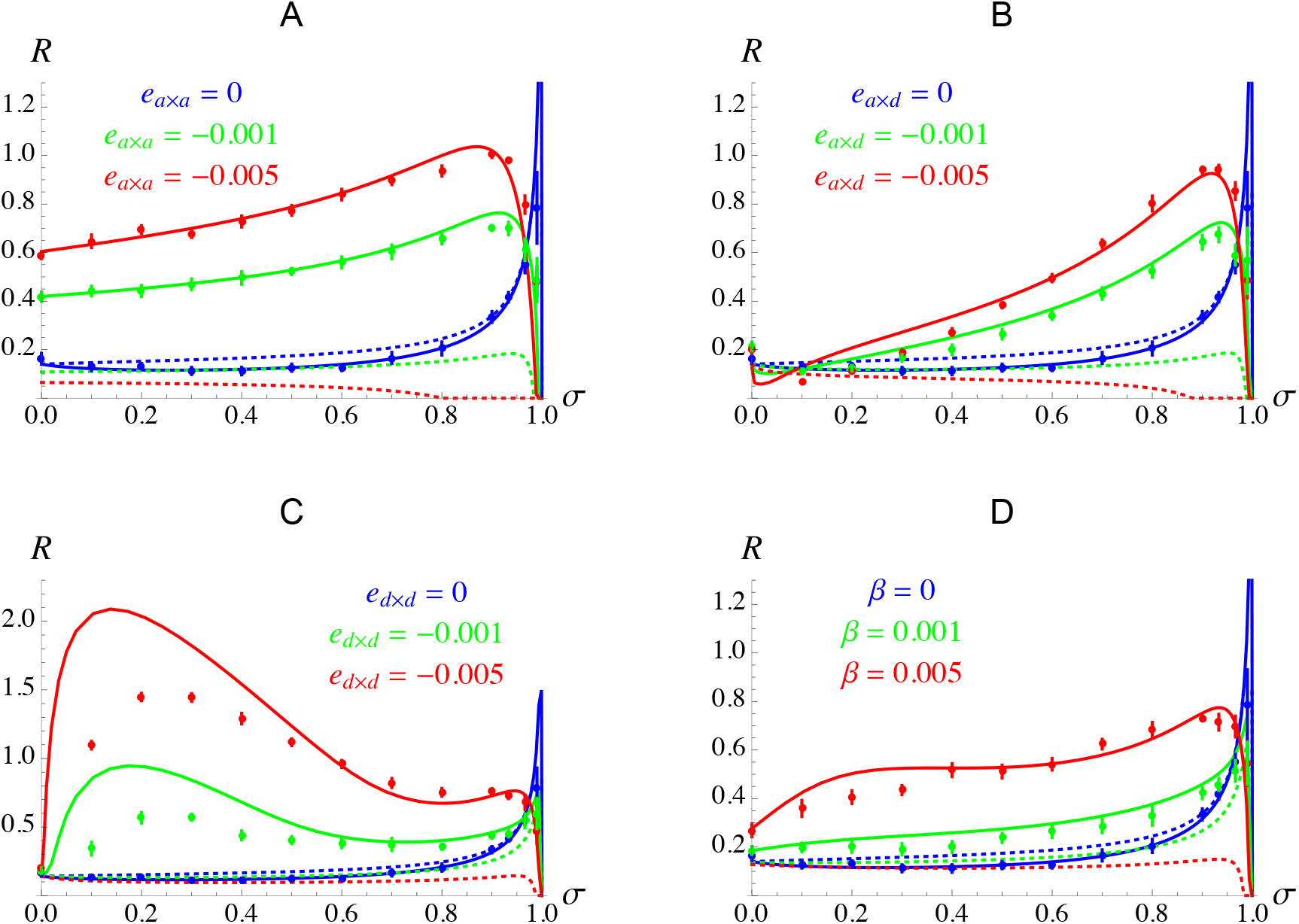
Effect of the selfing rate *σ* on the evolutionarily stable map length *R*_ES_, for different values of additive-by-additive (A), additive-by-dominance (B) and dominance-by-dominance (C) epistasis, and different values of Charlesworth et al.’s (1991) synergistic epistasis coefficient *β* (D). Note the different scale of the y-axis in C. Dots correspond to simulation results, curves have the same meaning as in Figure 3, and default parameter values are as in Figure 3 with *c* = 0.001.

## DISCUSSION

Digging into the causes of general empirical patterns such as the positive correlation between selfing rate and chiasma frequency observed within several families of flowering plants may help us to gain a better understanding of the selective forces affecting the evolution of recombination rates. While there is no obvious reason why the mechanistic constraints associated with chromosomal segregation during meiosis should differ between outcrossing and selfing species, the mating system of organisms does affect the benefit of recombination through its effect on genetic variation. Although indirect selective forces acting on recombination are expected to vanish under complete selfing (as heterozygosity should then be very rare), the results presented in this article show that intermediate selfing rates may either increase or decrease selection for recombination caused by interference (Hill-Robertson effect) among deleterious mutations, depending on parameter values. Roughly, selfing leads to stronger selection for increased chromosomal map length as long as *sh* ≪ *R* (1 – *F*) (mostly due to the fact that selfing reduces effective recombination rates, thus increasing the strength of genetic associations), while selfing decreases selection for recombination when *sh* » *R* (1 – *F*), due to the fact that changes in recombination rates have only little effect on genetic associations in this regime. Given that most deleterious mutations seem to have weak fitness effects (e.g., Charlesworth, 2015), selection for recombination should thus be increased by selfing, and be maximized for selfing rates slightly below 1 (as illustrated by Figures 3 and 4).

In agreement with previous results (Otto and Barton, 2001; Keightley and Otto, 2006; Roze, 2021), we found that interference is often the main driver of selection for recombination when linkage is tight (or when the selfing rate is strong), while deterministic effects tend to become more important when recombination is abundant. In the absence of epistasis, the variance in the degree of inbreeding among individuals caused by partial selfing (associated with a more efficient purging of deleterious alleles in more inbred lineages) generates positive associations among deleterious alleles, selecting against recombination in infinite populations. As a result, the equilibrium map length may be lower under moderate selfing rates than under random mating in large, highly recombining populations, as can be seen on Figure 3 with *c* = 10^−4^. As in standard models of infinite, randomly-mating populations (Charlesworth, 1990; Barton, 1995), negative epistasis between mutations tends to favor recombination by generating negative linkage disequilibria between deleterious alleles. This effect is often maximized at high selfing rates, again due to the fact that selfing reduces effective recombination rates, thus increasing the magnitude of linkage disequilibria. Furthermore, components of epistasis involving dominance (*e*_a×d_, *e*_d×d_) also contribute to generating linkage disequilibrium (through an effective epistasis parameter) when the selfing rate is greater than zero.

Under high effective recombination, Roze and Lenormand (2005) showed that correlations in homozygosity among loci generated by partial selfing (identity disequilibria) are the main drivers of indirect selection on recombination rates, favoring higher recombination when dominance-by-dominance epistasis is negative (as recombination then benefits from a short-term advantage, by increasing the mean fitness of offspring). The results of the present article tune down the relative importance of this effect, by showing that it becomes negligible relative to the effect of deterministically generated linkage disequilibria and Hill-Robertson interference under high selfing and in the case of tightly linked loci. Nevertheless, moderate selfing rates may favor a significant increase in chromosomal map length through this short-term benefit of breaking identity disequilibria, provided that dominance-by-dominance interactions are negative and represent a major component of epistasis (Figure 5C). In principle, the average sign and overall importance of dominance-by-dominance effects can be inferred from the shape of the relation between the degree of inbreeding of individuals and their fitness (Crow and Kimura (1970), p. 80), negative *e*_d_×_d_ causing a faster than linear decline in fitness with inbreeding. This method was used on several plant species and did not yield any clear evidence for negative *e*_d_×_d_ (Willis, 1993; Falconer and Mackay, 1996); however, the methodology used (involving experimental crosses to increase the degree of inbreeding of individuals) generates a bias against finding negative *e*_d×d_, as deleterious alleles may have been purged from the more highly inbred lines. Using an experimental protocol that avoids this bias, Sharp and Agrawal (2016) found evidence for negative e_d_×_d_ (on viability) between EMS-induced mutations in *Drosophila melanogaster*. While more work is needed to assess the generality of this result, previous experimental studies showed that epistasis is typically quite variable among pairs of loci (de Visser and Elena, 2007; Kouyos et al., 2007; Martin et al., 2007). As shown by Otto and Feldman (1997), recombination tends to be less favored when epistasis is variable, and it would thus be of interest to extend our model to more realistic fitness landscapes including distributions of epistasis.

While the indirect benefits of increased crossover rates may be strong when recombination is rare (e.g., Keightley and Otto, 2006), they typically become rather weak under frequent recombination (Roze, 2021), and one may thus wonder to what extent our model can explain a positive effect of selfing on chiasma frequency when at least one crossover per chromosome occurs during meiosis. When our model is modified so that 1 crossover per bivalent necessarily occurs (leading to a minimum map length of 50cM, that is, *R* = 0.5) and letting *R* evolve above 0.5 using a similar model as before, one indeed observes very limited effects of indirect selection on the evolutionarily stable map length (except for high selfing rates) in the absence of epistasis, for a deleterious mutation rate per chromosome of *U* = 0.1 (Figure 6). Substantial increases in recombination under partial selfing can be favored in the presence of negative *e*_d×d_, however, chromosomal map length being maximized for moderate selfing rates in this case (Figure 6A). Furthermore, higher chromosomal mutation rates lead to important increases in *R* at high selfing rates due to stronger Hill-Robertson effects, as can be seen on Figure 6B for *U* = 0.5. Relaxing the (probably unrealistic) assumption that crossovers occur anywhere along the chromosome with the same probability may also generate stronger effects of indirect selection on the evolutionary stable map length. In particular, data from *Caenorhabditis elegans* and humans suggest that structural constraints may impose restrictions on the possible localization of obligate crossovers along chromosomes (Koehler et al., 1996; Ottolini et al., 2015; Altendorfer et al., 2020); the fact that crossovers are often more frequent in subtelomeric regions (in plants and animals) may possibly reflect such constraints (e.g., Haenel et al., 2018). Restricting the position of the obligate crossover in a given chromosomal region should increase the strength of indirect selection acting on a modifier allele increasing recombination in other regions, particularly if the modifier is located in regions with lower recombination. Effects of non-uniform positions of crossovers along chromosomes and crossover interference will be explored in a future work. Sweeps of beneficial mutations may also increase the strength of indirect selection for recombination (Hartfield et al., 2010; Roze, 2021). In particular, transitions towards predominant selfing are often followed by a number of phenotypic changes such as reduced flower size (“selfing syndrome”, e.g., Cutter, 2019) that may be seen as adaptations to the new mating system, thus involving the spread of newly beneficial alleles that may (at least transiently) favor higher recombination rates. Recombination rates may also be further increased after transitions to selfing due to stronger Hill-Robertson effects caused by population bottlenecks and extinction/recolonization dynamics that often characterize selfing populations (e.g., Guo et al., 2009; Willi et al., 2018; Orsucci et al., 2020).

**Figure 6.**
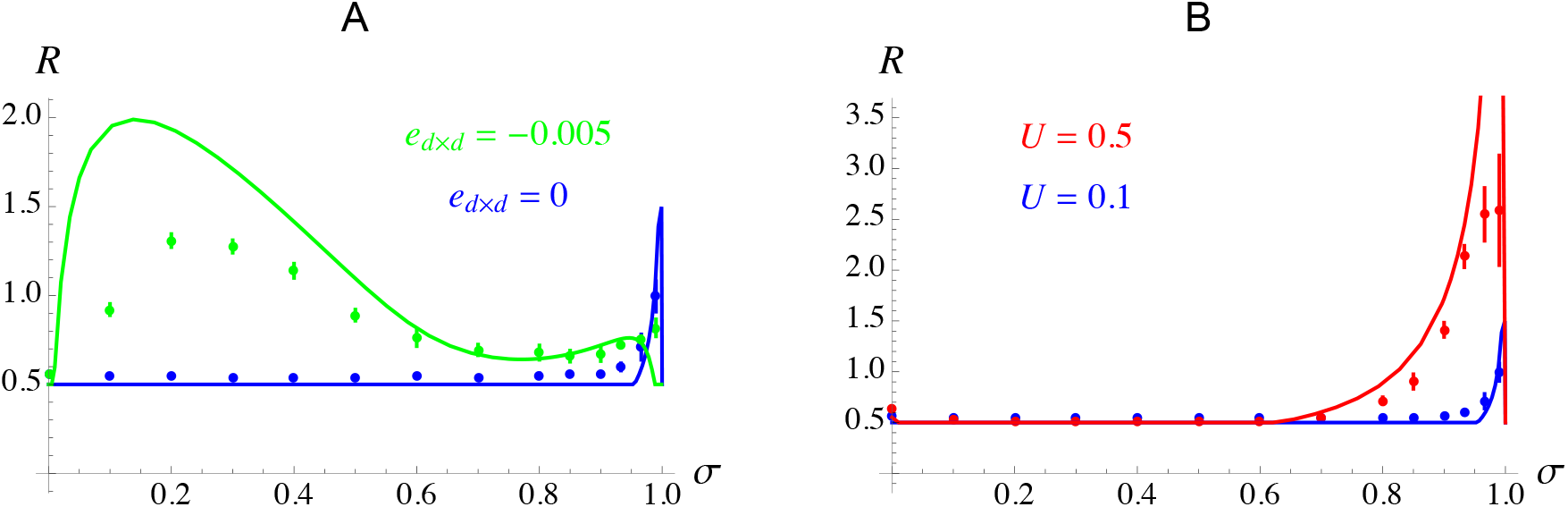
Effect of the selfing rate *σ* on the evolutionarily stable map length *R*_ES_, when at least one crossover per chromosome occurs during meiosis (leading to a minimal map length of *R* = 0.5. Dots correspond to simulation results, curves correspond to analytical predictions; default parameter values are as in Figure 3 with *c* = 0.001. A: Blue: no epistasis, green: *e*_d×d_ = –0.005. B: Blue: *U* = 0.1, red: *U* = 0.5.

Interestingly, predominantly selfing species often maintain low rates of outcrossing in nature (e.g., Bonnin et al., 2001; Bomblies et al., 2010), which may possibly be the consequence of selective forces favoring the maintenance of recombination. Indeed, deterministic and stochastic multilocus models showed that individuals outcrossing at a low rate are often selectively favored over complete selfers in conditions where recombination is advantageous (negative epistasis or finite population size, Charlesworth et al., 1991; Kamran-Disfani and Agrawal, 2014). However, higher effective recombination rates could in principle be achieved by increasing either the outcrossing rate of individuals or the number of crossovers at meiosis, and one could imagine that one solution or the other may be favored depending on the different types of direct and indirect selective forces that may act on outcrossing vs. recombination modifiers. Exploring the joint evolution of outcrossing and recombination rates in predominantly selfing species would thus be of interest, both from a theoretical and empirical perspective.

## Acknowledgements

We thank the bioinformatics and computing service of Roscoff’s Biological Station (Abims platform) for computing time, and the Agence Nationale pour la Recherche for funding (SelfRecomb project: ANR-18-CE02-0017-02, and GenAsex project: ANR-17-CE02-0016-01).

## APPENDIX A: HIGH EFFECTIVE RECOMBINATION

Under high effective recombination, the associations between the modifier and the selected loci involved in equation 6 are given by (see *Mathematica* notebook for derivation):

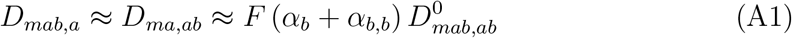

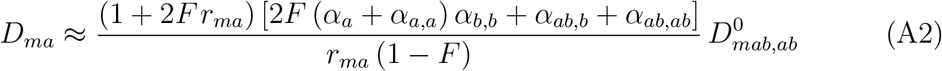

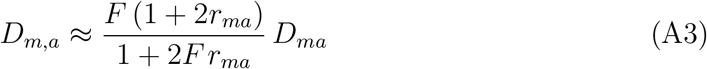

*D_mab,b_, D_mb,ab_, D_mb_* and *D_m,b_* being given by symmetric expressions. *D_mab_, D_m,ab_* and *D_ma,b_* are given by:

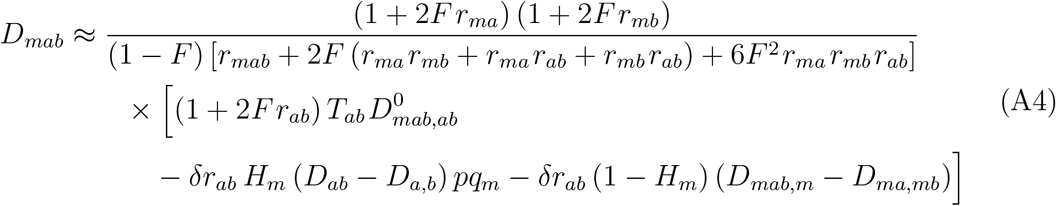

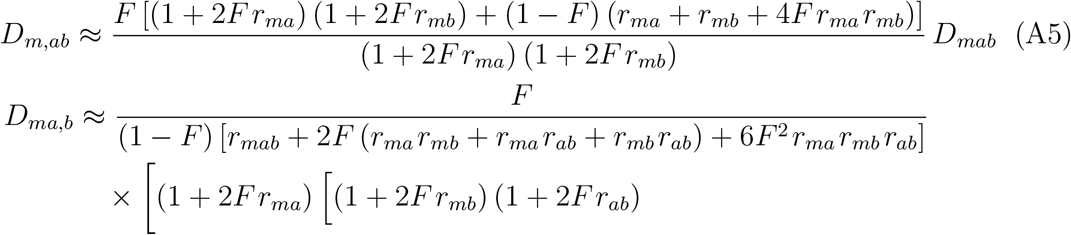

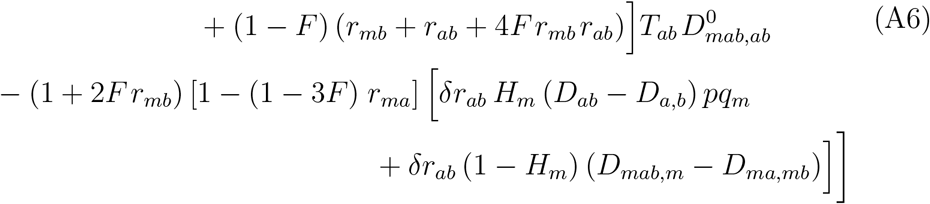

(*D_mb,a_* being given by a symmetric expression), with *H_m_* = *h_m_* + (1 – 2*h_m_*)*p_m_*, and:

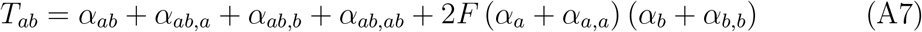

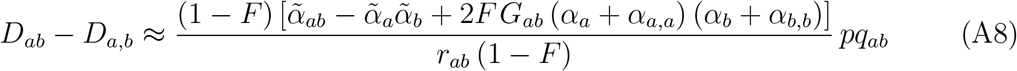

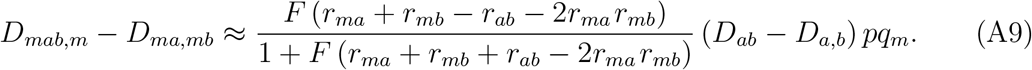

The identity disequilibrium *G_ab_* is given by *ϕ_ab_* – *F*^2^, with:

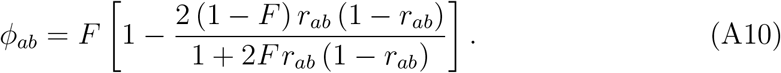

## APPENDIX B: INTERPRETING EFFECTIVE SELECTION COEFFICIENTS

The form of the coefficients 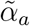 and 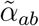 representing the effective strength of selection against deleterious alleles and the effective epistasis between pairs of deleterious alleles in a partially selfing population can be understood as follows. Assuming that deleterious alleles stay rare in the population (*u* ≪ *s*), we can neglect homozygotes for deleterious alleles produced by outcrossing, and consider that homozygosity for deleterious alleles necessarily implies identity-by-descent among these alleles. Then, if an individual carries allele *a* on one of its haplotypes, with probability *F* it also carries allele *a* on its second haplotype. In that case, the fitness effect of the deleterious allele (*α_α_*) is increased by *α_a_* due to the presence of *a* on the second haplotype, and by *α_a,a_* due to the interaction among those two deleterious alleles: therefore, the effective strength of selection experienced by deleterious alleles is 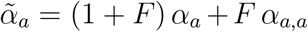 (which, using equation 5, yields 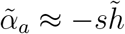). A similar reasoning can be used to compute the effective epistasis coefficient. With probability *ϕ_ab_*, an individual is identical-by-descent at both loci *a* and *b*; with probability *F* – *ϕ_ab_* it is identical-by-descent at the first locus and not at the second (or at the second and not at the first), while with the complementary probability 1 – 2*F* + *ϕ_ab_* it is identical-by-descent at neither locus. Therefore, an individual carrying alleles *a* and *b* on one of its haplotypes is *aabb* with probability *ϕ_ab_, aaBb* (or *Aabb*) with probability *F* – *ϕ_ab_*, and *AaBb* with probability 1 + 2*F* – *ϕ_ab_*. In the first case (*aabb*), the fitness effect of the interaction between *a* and *b* (*α_ab_*) is increased by the interaction between *a* and *b* present on the other haplotype (*α_ab_*), twice the interaction between deleterious alleles in trans (*α_a,b_*), twice the additive-by-dominance interactions (*α_ab,a_, α_ab,b_*), and by the dominance-by-dominance interaction (*α_ab,ab_*). In the second case (*aaAb* or *Aabb*), it is increased by the interaction between deleterious alleles in trans (*α_a,b_*) and by the additive-by-dominance interaction (*α_ab,a_* or *α_ab,b_*). This yields:

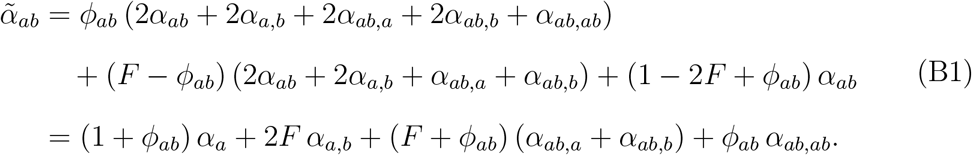

## APPENDIX C: GENERAL QLE APPROXIMATIONS

The following expressions combine approximations obtained under different regimes (high effective recombination, weak recombination, high selfing, epistasis of order *ϵ*^2^ or *ϵ*, see Supplementary Material). The different associations affecting the change in frequency of the modifier (equation 6) are given by (with 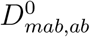 given by equation 10):

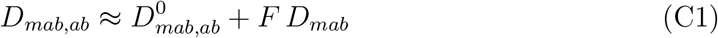

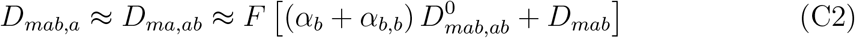

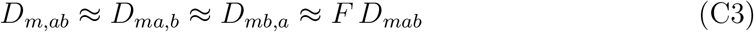

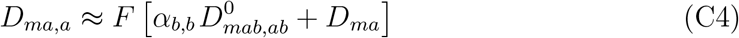

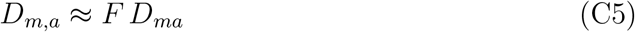

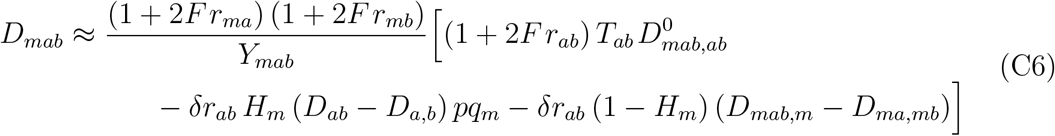

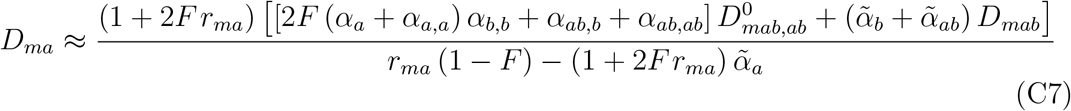

(*D_mab,b_, D_mb,ab_, D_mb,b_, D_m,b_* and *D_mb_* being given by symmetric expressions), with:

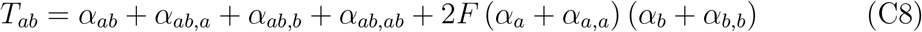

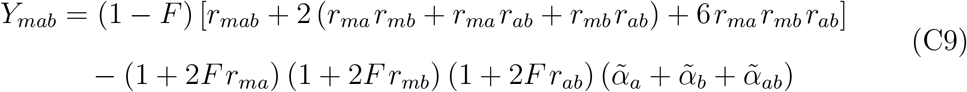

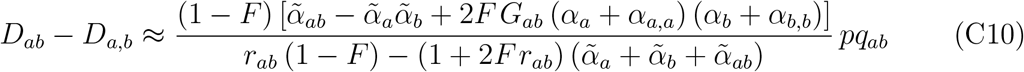

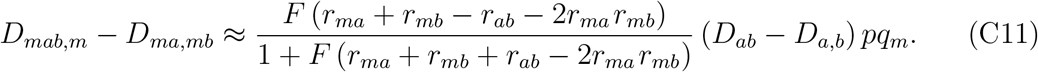

The change in frequency of the modifier is given by 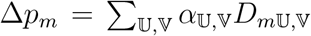, where the double sum is over all elements of the set {Ϙ, *a, b, ab*}, and thus decomposes into a term generated by 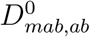 (that becomes predominant under high effective recombination) and a term generated by *D_ab_* – *D_a,b_* (that becomes predominant under weak recombination or strong selfing).

