## Supplementary Figures for "The evolution of recombination in self-fertilizing organisms"

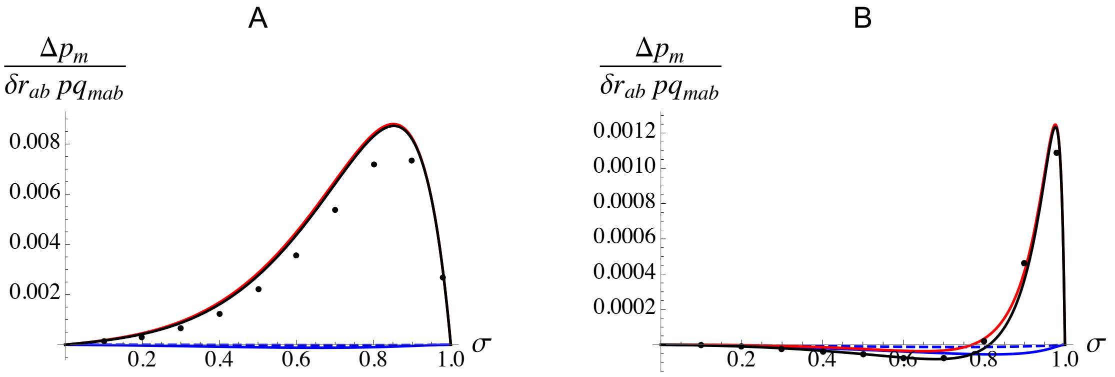


**Figure S1.** Same as Figure 1 with $e_{\mathrm{a}\times\mathrm{d}} = -0.001$, and $r_{ma} = r_{ab} = 0.01$ (A), $r_{ma} = r_{ab} = 0.1$ (B). Other parameter values are as in Figure 1.

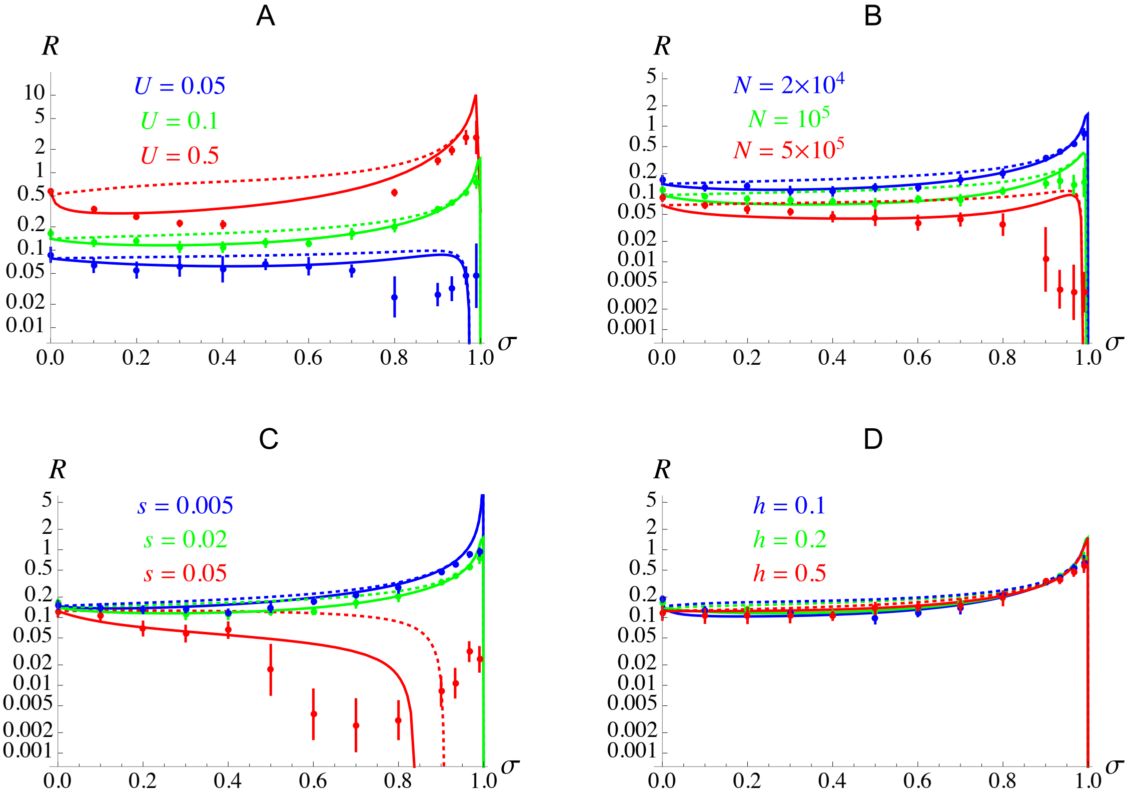


**Figure S2.** Same as Figure 4, with $R$ on a log scale.

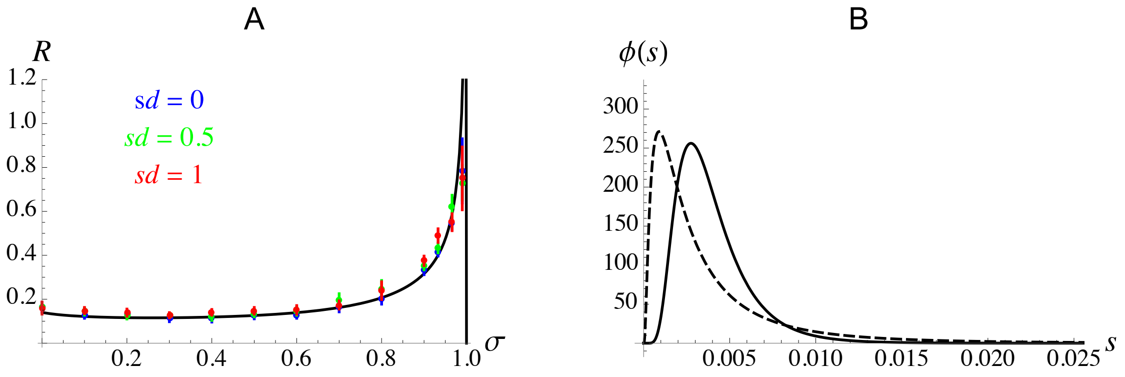


**Figure S3.** A: effect of the selfing rate $\sigma$ on the evolutionarily stable map length $R_{\mathrm{ES}}$ for different values of the standard deviation $sd$ of the log-normal distribution of selection coefficients $s$ of deleterious alleles (in the simulations). B: distribution of $s$ for $sd = 0.5$ (solid) and $sd = 1$ (dashed). The mean selection coefficient $\overline{s}$ is set at 0.02, and default parameter values are as in Figure 3 with $c = 0.001$. The analytical prediction (curve) in A is the same as in Figure 3 for $c = 0.001$ (fixed $s$).

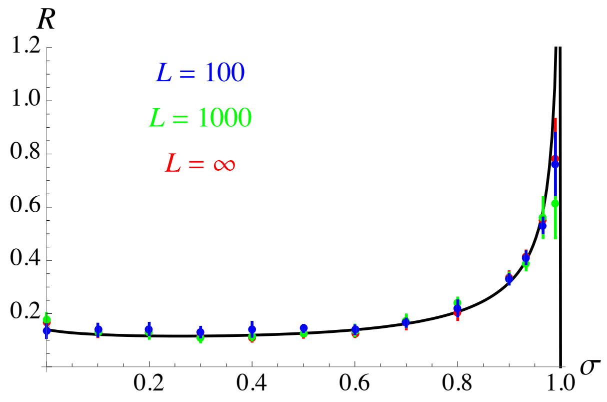


**Figure S4.** Effect of the selfing rate $\sigma$ on the evolutionarily stable map length $R_{\mathrm{ES}}$ for different numbers of loci $L$ at which deleterious alleles may occur. In simulations with finite $L$, loci are uniformly spaced along the chromosome, each locus mutating at a rate $U/L$. Parameter values are as in Figure 3 with $c = 0.001$. The analytical prediction (curve) is the same as in Figure 3 for $c = 0.001$ (infinite number of loci).

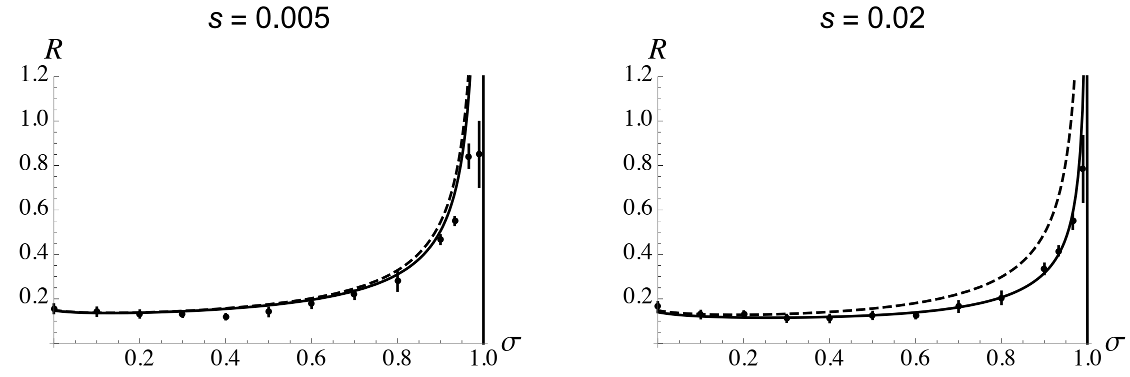


**Figure S5.** Effect of the selfing rate $\sigma$ on the evolutionarily stable map length $R_{\mathrm{ES}}$ for $s = 0.005$ and $s = 0.02$, and other parameter values as in Figure 4. Solid curves are the same as in Figure 4C, corresponding to the predictions obtained by solving $s_{\mathrm{direct}} + s_{\mathrm{det}} + s_{\mathrm{HR}} = 0$ for $R$, where $s_{\mathrm{det}}$ and $s_{\mathrm{HR}}$ are obtained by integrating the expressions for the strength of indirect selection generated by deterministic effects (for $s_{\mathrm{det}}$) and by the Hill-Robertson effect (for $s_{\mathrm{HR}}$) over the genetic map (see Supplementary Material). Dashed curves correspond to the predictions obtained using the same method, but replacing $s_{\mathrm{HR}}$ by the approximation given by equation 22: $s_{\mathrm{HR}} \approx 1.8\delta R\, U^2 / \left[N_{\mathrm{e}} R^3 \left(1 - F\right)^2\right]$.

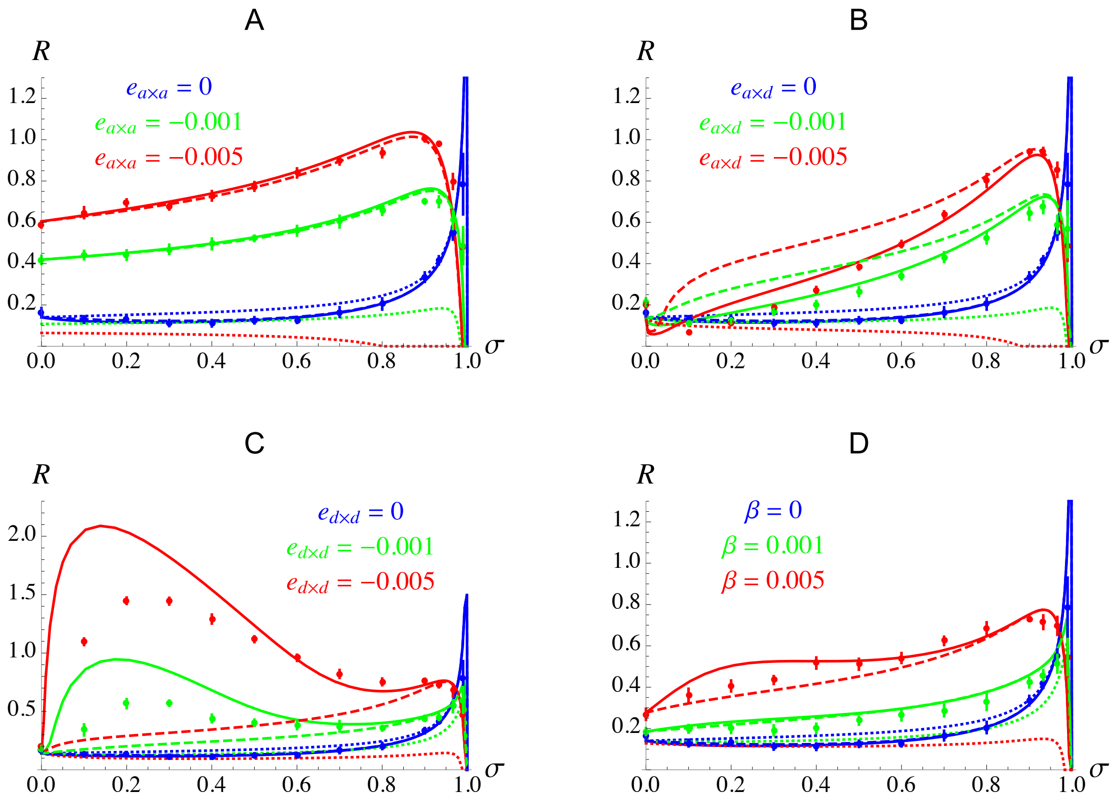


**Figure S6.** Same as Figure 5, the dashed curves showing the predictions obtained for $R_{\mathrm{ES}}$ when ignoring the term generated by $D^0_{mab,ab}$ in the expression for the deterministic source of indirect selection acting on the modifier locus. The difference between the dotted and dashed curves therefore corresponds to the effect of deterministic terms generated by $D_{ab}$, while the difference between the dashed and solid curves corresponds to the effect of deterministic terms generated by $D^0_{mab,ab}$.
